# Digital control of c-di-GMP in *E. coli* balances population-wide developmental transitions and phage sensitivity

**DOI:** 10.1101/2021.10.01.462762

**Authors:** Alberto Reinders, Benjamin Sellner, Firas Fadel, Margo van Berkum, Andreas Kaczmarczyk, Shogo Ozaki, Johanna Rueher, Pablo Manfredi, Matteo Sangermani, Alexander Harms, Camilo Perez, Tilman Schirmer, Urs Jenal

## Abstract

Nucleotide-based signaling molecules (NSMs) are widespread in bacteria and eukaryotes, where they control important physiological and behavioral processes. In bacteria, NSM-based regulatory networks are highly complex, entailing large numbers of enzymes involved in the synthesis and degradation of active signaling molecules. How the converging input from multiple enzymes is transformed into robust and unambiguous cellular responses has remained unclear. Here we show that *Escherichia coli* converts dynamic changes of c-di-GMP into discrete binary signaling states, thereby generating heterogeneous populations with either high or low c-di-GMP. This is mediated by an ultrasensitive switch protein, PdeL, which senses the prevailing cellular concentration of the signaling molecule and couples this information to c-di-GMP degradation and transcription feedback boosting its own expression. We demonstrate that PdeL acts as a digital filter that facilitates precise developmental transitions, confers cellular memory, and generates functional heterogeneity in bacterial populations to evade phage predation. Based on our findings, we propose that bacteria apply ultrasensitive regulatory switches to convert dynamic changes of NSMs into binary signaling modes to allow robust decision-making and bet-hedging for improved overall population fitness.

## Introduction

Biological systems need to convert spatial or temporal gradients of signaling molecules into precise and robust readouts. For example, in the *Drosophila* embryo, highly accurate spatial patterning is established from an original gradient of the maternal morphogen *bicoid* within the first few hours of development (Gregor et al., 2007; Jaeger, 2011). Gradual changes of *bicoid* concentrations along the embryonic axis are converted into sharp expression patterns of its downstream target gene *hunchback* via switch-like, cooperative responses (Driever et al., 1989; Park et al., 2019). Similar mechanisms must exist to convert gradual changes of signaling molecules into specific and robust cellular responses. This is of particular relevance for bacteria, which make use of an extensive array of sensory systems for surveillance. These include receptors coupled to protein phosphorylation cascades (Bi and Sourjik, 2018; Capra and Laub, 2012; Galinier and Deutscher, 2017) or to nucleotide-based signaling molecules (NSM) (Bassler et al., 2018; Hauryliuk et al., 2015; Jenal et al., 2017; Stülke and Krüger, 2020; Zaver and Woodward, 2020). While phosphorylation cascades are generally linear and highly specific, networks relying on small signaling molecules are less well defined.

NSMs are widespread in bacteria and include the linear (p)ppGpp, monocyclic compounds like cAMP, and cyclic di-nucleotides like c-di-GMP, c-di-AMP, or cGAMP. These molecules control core bacterial processes like growth and metabolism, stress response and predator defense, as well as virulence and persistence (Bassler et al., 2018; Hauryliuk et al., 2015; Jenal et al., 2017; Stülke and Krüger, 2020; Zaver and Woodward, 2020). NSM-mediated signaling networks can adopt highly complex architectures with many bacteria harboring up to several dozens of sensors regulating the concentration of active compounds. For example, some spirochetes, alpha-proteobacteria, or mycobacteria possesses up to 30 adenylate cyclases, while many actinobacteria, cyanobacteria, or proteobacteria encode 50-100 different enzymes involved in the synthesis and breakdown of c-di-GMP (Bassler et al., 2018; Galperin, 2010, 2018).

This raises important questions regarding NSM control and downstream signaling. How do bacteria convert gradual temporal changes of signaling molecules into deterministic and irreversible cellular responses? And how do they absorb stochastic fluctuations of such molecules generated through random noise? This is particularly important if downstream processes are highly sensitive to concentration changes and if they occur on short time scales. For example, effector binding affinities for c-di-GMP are typically in the nanomolar range (Chou and Galperin, 2016) and some c-di-GMP-mediated processes show ultra-rapid responses on the time scale of seconds (Hug et al., 2017; Laventie et al., 2019; Nesper et al., 2017).

Finally, how can small diffusible molecules regulate specific downstream processes given that concentration changes likely provoke a global cellular response? It was recently proposed that c-di-GMP can signal in spatially confined compartments with specialized sensors stimulating spatially coupled cellular processes (Lori et al., 2015; Richter et al., 2020). Although ‘local signaling’ relies on a direct interaction between sensors regulating c-di-GMP synthesis and/or degradation and its downstream targets (Andrade et al., 2006; Dahlstrom et al., 2016; Giacalone et al., 2018; Lindenberg et al., 2013), molecule leakage likely occurs and needs to be absorbed to effectively isolate individual signaling modules from each other (Jenal et al., 2017; Richter et al., 2020).

One possibility to avoid detrimental fluctuations of potent signaling molecules and to convert graded inputs into switch-like, irreversible and specific responses is to sense the prevailing concentration and couple this information to catalytic feedback. Positive and double-negative feedback loops can generate stable genetic responses when coupled to non-linear or ‘ultrasensitive’ behavior (Ferrell, 2002; Koshland et al., 1982). In recent years, several examples of switches were identified generating bi-stable gene expression and cell fate decisions in bacteria (Losick and Desplan, 2008; Norman et al., 2015; Veening et al., 2008). However, to date, no such mechanisms are known to control the concentration of small diffusible signaling molecules like cAMP or c-di-GMP.

Here, we show that *Escherichia coli* converts gradual changes of c-di-GMP into a discrete binary output, thereby generating heterogeneous populations with either high or low c-di-GMP levels. This response is mediated by the phosphodiesterase PdeL that acts as a simple molecular switch establishing precise cellular levels of c-di-GMP. Our results show that PdeL degrades c-di-GMP and, at the same time, acts as a c-di-GMP-dependent transcription factor to regulate its own expression. We demonstrate that c-di-GMP impedes both PdeL activities and that catalytic and transcriptional feedbacks of PdeL generate bimodal and bistable populations with distinct behavior. Functional assays indicate that the PdeL switch sets off robust lifestyle changes that lead to precise biofilm formation and efficient escape and that it can serve as bet-hedging device to protect *E. coli* against phage predation.

## Results

### Transcription of the *pdeL* phosphodiesterase gene is controlled by c-di-GMP

Earlier work had shown that PdeL autoregulates its own transcription (Reinders et al., 2015). To dissect PdeL transcription control, DNA shift assays were carried out with fragments spanning the entire *pdeL* promoter region (Shimada et al., 2005). This identified two binding sites for PdeL upstream of the promoter, a more distal one and one closer to the promoter, which overlaps with the binding site for the central metabolic regulator Cra (Kochanowski et al., 2013; Shimada et al., 2011) **(Fig. 1a,b)**. Both Cra (K_d_ 49 nM) and PdeL (K_d_ 76 nM) bind to the proximal site with high affinity **(Fig. S1a-d)**. While Cra binding is independent of PdeL, PdeL binding requires Cra **(Fig. 1b,c)**. Hence, we termed the proximal PdeL binding motif the Cra-dependent PdeL-box (CDB) to distinguish it from the upstream Cra-independent binding site (CIB), for which PdeL showed significantly lower affinity (K_d_ 573 nM) **(Figs. 1b,c; S1e,f)**.

**Figure 1:**
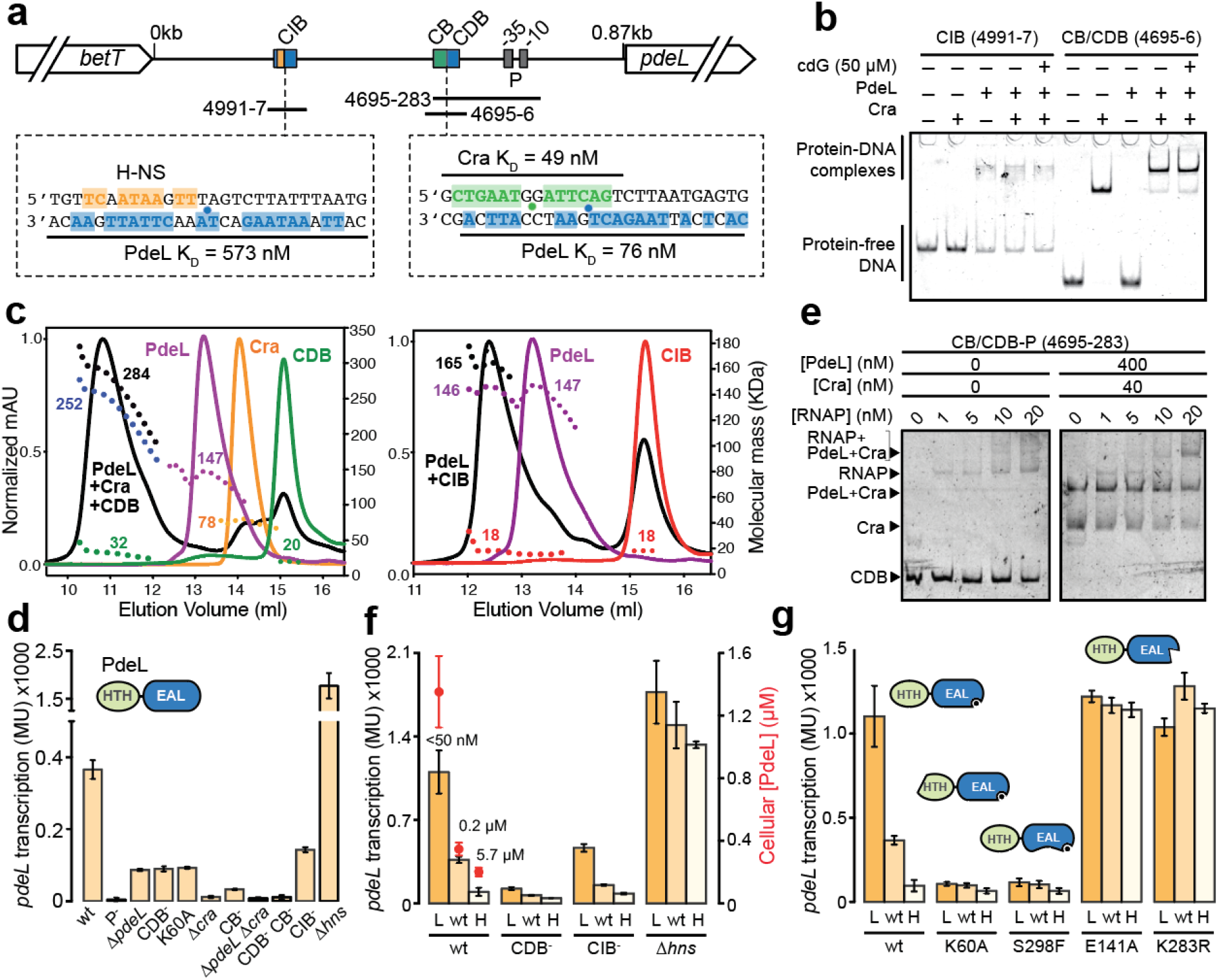
Regulation of *pdeL* transcription. **(a)** Schematic of *pdeL* promoter region. Binding sites for PdeL (CDB = **<u>C</u>**ra-**<u>d</u>**ependent PdeL-**<u>b</u>**ox; CIB = **<u>C</u>**ra-**i**ndependent PdeL-**<u>b</u>**ox), Cra (CB) and H-NS are shown in blue, green and orange, respectively. Palindromic residues of the CIB and homologous residues in the CDB are highlighted in blue and are shown on the lower strand; residues of the H-NS and Cra binding sites are shown on the upper strand. Blue and green dots mark the center of palindrome sites. Binding affinities of PdeL and Cra are indicated. H-NS binding site is taken from the Virtual Footprint website http://www.prodoric.de/vfp/ (Münch et al., 2005). **(b)** Electrophoretic mobility shift assay (EMSA) of 5’ Cy3 labeled oligonucleotides and purified Cra- StrepII and PdeL-StrepII proteins. The position of the labeled oligonucleotides in the *pdeL* promoter region is indicated in (a). **(c)** SEC-MALS analysis of PdeL, Cra, and double-stranded DNA fragments (Left: CDB; right: CIB) containing the Cra and/or PdeL binding sites (see: (a)). The molecular masses of individual components and respective DNA-protein complexes are indicated. **(d)** Activity of *pdeL* promoter in *E. coli* wild type and different mutant strains as indicated. CIB^-^, CDB^-^ and CB^-^: sequences of binding sites were randomized to abolish TF binding (see: Fig. S1g). Deletion mutants and a point mutation abolishing DNA binding of PdeL (K60A) are indicated. Inset shows domain architecture of PdeL with DNA binding domain (HTH) in green and catalytic EAL domain in blue. **(e)** Recruitment of RNA polymerase (RNAP) to the *pdeL* promoter. Increasing concentrations of RNAP were incubated with 5’ Cy3 labeled target DNA (4695-283; see: (a)) in the absence or presence of 40 nM Cra-StrepII and 400 nM PdeL-StrepII. **(f)** Effect of c-di-GMP on *pdeL* transcription. *pdeL* transcription was assayed in strains with different c-di-GMP concentrations (L = **l**ow; wt; H = **<u>h</u>**igh). Levels of c-di-GMP are shown, and cellular concentrations of PdeL are indicated in red above the corresponding bars. **(g)** Transcription of *pdeL* in strains with different c-di-GMP concentrations. *pdeL* mutant alleles included K60A (DNA-binding), S298F (dimerization), E141A (c-di-GMP binding) and K283R (catalytic base).

Transcription of *pdeL* was strongly reduced in strains lacking Cra or PdeL and in strains with scrambled binding sites for Cra or PdeL **(Fig. 1d, S1g)**. To investigate how PdeL stimulates its own transcription, we analyzed the binding of RNA polymerase (RNAP) to the *pdeL* promoter. While RNAP alone showed poor interaction with the *pdeL* promoter region, interaction was strongly increased in the presence of Cra and PdeL **(Fig. 1e)**, arguing that PdeL facilitates RNAP recruitment. In addition to PdeL and Cra binding motifs, the *pdeL* promoter region contains several binding sites for the transcriptional silencer H-NS (Rangarajan and Schnetz, 2018) **(Fig. S1g)**, one of which overlaps with the left half-site of the CIB palindrome **(Fig. 1a)**. H-NS and PdeL compete for this site *in vitro* **(Fig. S1h)** and *pdeL* transcription was strongly derepressed in an *hns* mutant **(Fig. 1d)**, arguing that PdeL stimulates *pdeL* transcription by acting as an anti-silencer. Together these experiments identified Cra and PdeL as co-activators of *pdeL* transcription and proposed that Cra facilitates PdeL binding, which in turn helps recruiting RNAP to the promoter region.

Previous work suggested that PdeL autoregulation is mediated by the cellular concentration of c-di-GMP (Reinders et al., 2015). Because c-di-GMP did not alter PdeL binding to CDB or CIB *in vitro* (**Fig. 1b**), we next monitored *pdeL* transcription *in vivo* in response to changes in c-di-GMP concentration. To tune cellular c-di-GMP levels, we made use of the observation that *E. coli* strain MG1655 (CGSC 7740) lacking the phosphodiesterase PdeH has strongly increased levels of c-di-GMP (Reinders et al., 2015). By replacing the chromosomal copy of *pdeH* with a plasmid-born and IPTG-inducible *pdeH*, we showed that c-di-GMP levels could be tuned from low nanomolar concentrations to above 5 µM, while a *pdeH* wild-type strain showed intermediate c-di-GMP concentrations (210 nM) **(Fig. 1f)**. The expression of a *pdeL-lacZ* reporter (Reinders et al., 2015) was maximal at low c-di-GMP concentrations but strongly declined at higher c-di-GMP levels. Accordingly, PdeL protein levels inversely scaled with c-di-GMP levels, ranging from 200 nM to 1.35 µM **(Fig. 1f)**.

C-di-GMP-dependent control of *pdeL* transcription strictly depends on CDB and to a lesser extent on CIB, arguing that c-di-GMP impacts *pdeL* transcription via PdeL binding to both sites. A mutant lacking H-NS largely abolished the effect of c-di-GMP on *pdeL* expression **(Fig. 1f)**.

To dissect the mechanism of c-di-GMP-dependent regulation, we analyzed *pdeL* transcription in strains expressing different PdeL variants. Mutations interfering with DNA binding (K60A) (Reinders et al., 2015) or PdeL dimerization (S298F) (Sundriyal et al., 2014) showed baseline *pdeL* promoter activity irrespective of c-di-GMP levels, *i*.*e*., mirroring *pdeL* promoter activity at high c-di-GMP. In contrast, substitutions of the conserved active site glutamate (E141), which is involved in c-di-GMP binding (Sundriyal et al., 2014), or the catalytic base residue (K283), which is strictly required for catalysis (Rao et al., 2008), resulted in complete de-repression of *pdeL* transcription even at high c-di-GMP concentrations **(Fig. 1g)**. Based on these findings, we propose that PdeL senses the prevailing c-di-GMP concentration and that it acts as a transcription factor to boost its own expression when cellular c-di-GMP levels are low.

### An active PdeL tetramer drives transcription in the absence of c-di-GMP

We have shown above that PdeL binds to two independent motifs upstream of the *pdeL* promoter, indicating that a higher order complex may be involved in transcription autoregulation. Thus, to better understand the molecular mechanism of c-di-GMP dependent transcription, we analyzed the full-length PdeL protein *in vitro*. PdeL harbors an N-terminal DNA binding domain (HTH) and a C-terminal catalytic domain (EAL). Earlier studies had shown that the isolated EAL domain displays a monomer - dimer equilibrium with a K_d_ of about 1 μM (Sundriyal et al., 2014). In contrast, mass photometry analyses revealed that PdeL is a dimer even at very low protein concentrations (25 nM) (**Fig. S2a**), indicating that the HTH and EAL domains synergistically stabilize the dimer conformation. Moreover, SEC-MALS studies demonstrated that PdeL is in a dynamic dimer-tetramer equilibrium (**Fig. 2a**). This was supported by microscale thermophoresis (MST) experiments that revealed a tetramerization K_d_ of 9 μM and a Hill coefficient of 1.4 **(Fig. S2b)**. Redistribution of fluorophore labeled PdeL was slow with an equilibrium being reached only after about 2 hours (**Fig. 2b**). This indicated that the PdeL tetramerization process is complex and potentially cooperative.

**Figure 2:**
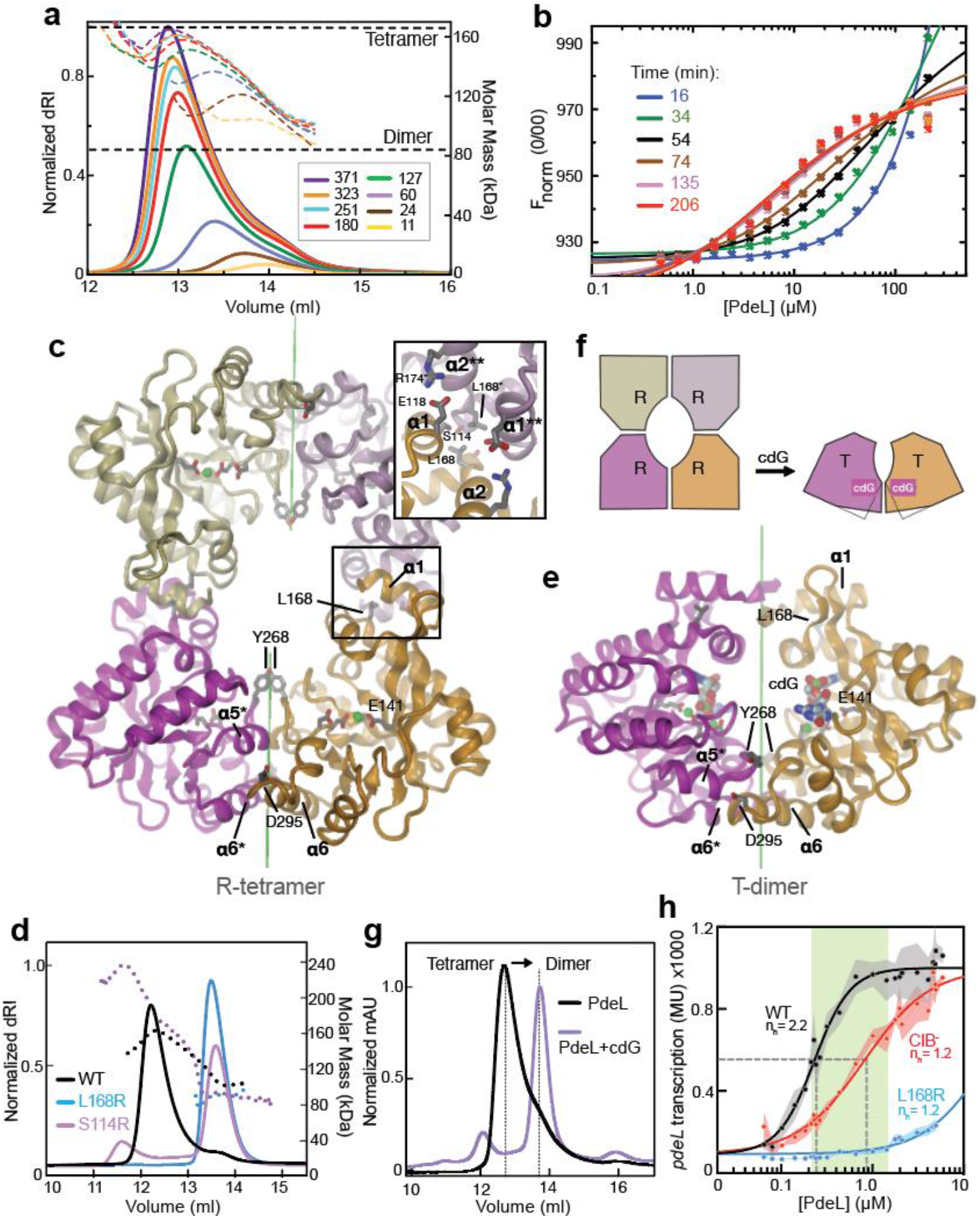
PdeL forms an active tetramer in the absence of c-di-GMP. **(a)** SEC-MALS analysis of PdeL at various loading concentrations (µM) as indicated. Full lines correspond to the normalized refractive index (dRI), stippled lines to molar mass (theoretical dimer and tetramer mass values are indicated). **(b)** MST analysis of PdeL dimer-tetramer equilibrium. PdeL dimer/tetramer exchange was measured by thermophoretic mobility upon titration of different amounts of purified Strep-tagged PdeL as indicated (µM) to a constant pool (50 nM) of fluorescently labeled His-tagged PdeL (PdeL*). Measurements were carried out repeatedly in roughly 18 min intervals as indicated until reaching equilibrium after 3.5 hrs. **(c)** Crystal structure of tetrameric PdeL. Tetrameric arrangement of PdeL EAL domains as seen in crystals of full-length PdeL (this study) and in PDB entry 4LYK. The two R-state dimers (with symmetry axes in green) are shown as cartoons, colored in bright and light magenta/orange, respectively. Selected residues are shown in full and are labeled, selected helices are labeled with asterisks denoting symmetry related helices. The inset shows details of the dimer-dimer contact. **(d)** SEC-MALS data of PdeL wild type and tetramerizaton mutants acquired at a loading concentration of 325 μM. **(e)** T-state dimer of the EAL domain of PdeL in complex with c-di-GMP (PDB: 4LJ3). **(f)** Schematic of the EAL domain arrangement of PdeL shown in **(c)** and **(e)** and proposed to be induced by c-di-GMP binding. Ligand binding is accompanied by a change in the intra-dimer interface resulting in a relative tilt of the monomers leading to tetramer dissociation. **(g)** SEC analysis of PdeL (120 μM) in absence and presence of c-di-GMP. **(h)** PdeL-dependent *pdeL* transcription in absence of c-di-GMP inhibition. Strains carried an IPTG inducible copy of *pdeH* on a plasmid to reduce c-di-GMP below the detection limit (see: Fig. 1f). To tune PdeL concentrations, *pdeL* was expressed from a tetracycline-inducible promoter (P_tet_-*pdeL*) (see: Fig. S3d,e and Materials and Methods for details). *lacZ* reporter fusions were used to assay *pdeL* promoter activity in wild type (black), CIB^-^ (red) and L168R mutants (tetramerization) (blue). Data points were fitted with an allosteric sigmoidal curve. Kinetic values were calculated for wild type: half maximal activation concentration, Khalf = 231 nM and n_h_ = 2.2; for CIB^-^: K_half_ = 800 nM and n_h_ = 1.2; and for L168R: K_half_ = 18 µM and h = 1.2. The green area depicts physiologically relevant cellular PdeL concentrations (see **Fig. 1f**).

Crystallization of full-length PdeL yielded a crystal form with four molecules in the asymmetric unit diffracting to 4.4 Å (Table S1). Molecular replacement with the known structure of the EAL domain of PdeL (4LYK) (Sundriyal et al., 2014) gave a clear solution for a tetramer with 222-symmetry (**Fig. 2c**). Re-analysis of the crystal structure of the EAL domain of PdeL showed that virtually the same tetrameric arrangement is present in the crystal lattice of 4LYK, suggesting that the tetramer assembly is not governed by crystal contacts. No solution was found for the HTH domains, indicating potential flexibility within the crystal lattice.

The tetramer is formed by head-to-head association of two canonical EAL dimers via two symmetric contact patches forming isologous, *i*.*e*., 2-fold symmetric interactions of helix α1 with α2 of the adjacent subunit (**Fig. 2c**). The dimer-dimer contacts are mediated by 14 individual residues with reciprocal salt bridges and hydrophobic residues contributing to the stabilization of the tetramer. Substitutions of S114 or L168, two central residues of the interface, rendered PdeL dimeric, as measured in solution at a concentration at which the wild-type proteins forms tetramers (**Fig. 2d**). Thus, the crystal structure likely represents the tetrameric species identified in solution. Intriguingly, based on steric reasons swapping of the HTH domains appears mandatory for the assembly of a full-length PdeL tetramer from two dimers (**Figs. S2c,d**), potentially explaining the observed slow tetramerization kinetics.

Structural analyses of the isolated EAL domain of PdeL had revealed two distinct dimer conformations, a canonical open (relaxed) state (R-state) and a closed (tight) dimer obtained in the presence of c-di-GMP (T-state) (Sundriyal et al., 2014). Intriguingly, the PdeL tetramer consists of two R-state dimers (**Fig. 2c**), while the closed T-state dimer **(Fig. 2e)** is unable to tetramerize due to the altered position and orientation of the dimer-dimer contact patches (**Fig. 2f**). In line with this, tetramerization of PdeL is abolished in presence of c-di-GMP as measured by SEC (**Fig. 2g**). To investigate the role of tetramerization in *pdeL* transcription, we monitored *pdeL* promoter activity at varying cellular PdeL concentrations in a strain expressing *pdeL* wild type or a tetramerization mutant from a tetracycline-inducible promoter (P_tet_*-pdeL*) **(Fig. S3a,b)**. To avoid interference from c-di-GMP, this strain also harbored an IPTG-inducible copy of *pdeH* (P_lac_*-pdeH*), which reduced c-di-GMP to below the limit of detection (50 nM) **(Fig. 1f)**. When PdeL concentrations were gradually increased, *pdeL* promoter activity increased in a highly cooperative manner (n_h_ = 2.2) with an activation constant of 230 nM PdeL **(Fig. 2h)**. In contrast, the tetramerization-deficient variant L168R completely abolished *pdeL* transcription at physiological PdeL concentrations **(Fig. 1f, 2h)**. Moreover, a mutant with a scrambled CIB binding site, although still induced at higher PdeL levels, showed a non-cooperative response (n_h_ = 1.2) (**Fig. 2h**). These data are in line with the observed PdeL binding affinity to CIB (K_D_ = 573 nM) and with the cellular concentrations of PdeL in the promoter OFF (200 nM) and ON states (1.35 µM) **(Figs. 1, S1)** and argue that binding of PdeL to CIB is required for the non-linear behavior of *pdeL* promoter activity.

Altogether, these findings propose that PdeL forms tetramers at low c-di-GMP concentrations and that this process is responsible for the cooperative, switch-like activation of *pdeL* transcription by engaging both CDB and CIB and possibly involving DNA looping to counteract H-NS silencing.

### Destabilizing the PdeL T-state conformation results in constitutive transcription activity

The above structural and biophysical experiments proposed a model in which c-di-GMP binding to PdeL mediates the thermodynamic interconversion between R-state tetramers and T-state dimers (**Fig. 2f**). To corroborate that PdeL changes its conformation in a c-di-GMP-dependent manner, we carried out cysteine cross-link experiments in the presence and absence of c-di-GMP. We chose to substitute Y268 with Cys because this residue is in close proximity with its symmetry mate in the R-state but distant in the T-state dimer (**Figs. 2c,e**). Indeed, oxidation of the Y268C mutant produced a cross-linked dimer, which was significantly reduced in presence of c-di-GMP (**Fig. 3a**), as expected for the T-state conformation.

**Figure 3:**
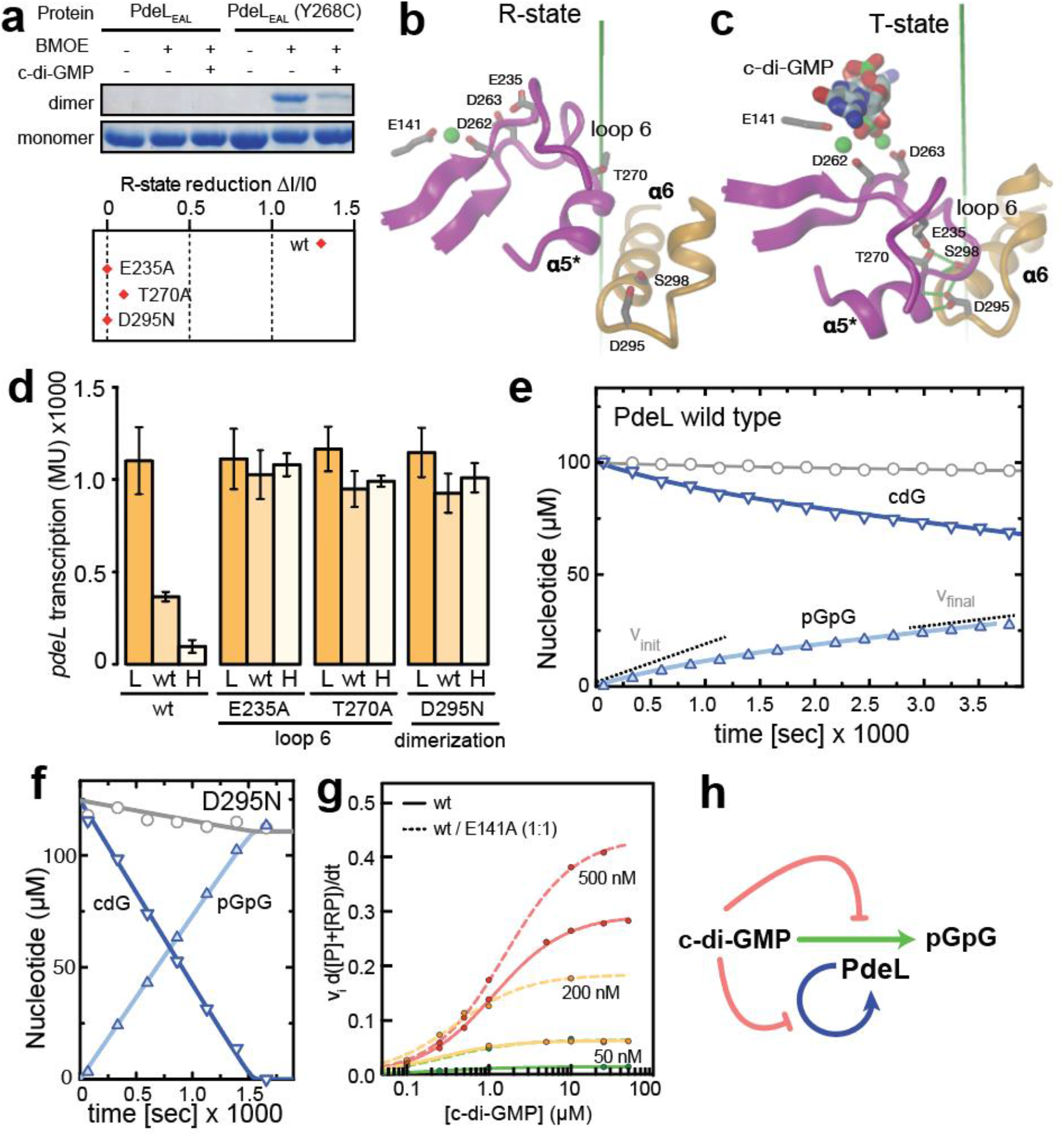
PdeL R-lock mutants show constitutive, c-di-GMP independent transcriptional and enhanced catalytic activity. **(a)** *In vitro* cysteine cross-linking of the PdeL Y268C mutant. Crosslinks were performed without and with 50 µM c-di-GMP and proteins were separated by SDS-PAGE. PdeL wild type is shown as negative control for crosslink specificity. Relative changes in c-di-GMP dependent crosslinking of PdeL wild type and R-lock mutants are shown in the graph below. R-state reduction of wild type and mutant proteins refers to the relative differences in abundance of crosslinked dimers as recorded in the absence and presence of c-di-GMP. **(b,c)** Structural details of intra-dimer interface of R-state **(b)** and T-state **(c)** of PdeL. Representations are as in Fig. 2. **(d)** *pdeL* promoter activity in strains carrying R-lock PdeL mutations (see panels a and b). L, wt, and H refer to levels of c-di-GMP as in Fig. 1f. **(e, f)** Enzymatic progress curves of wild-type PdeL **(e)** and R-lock mutant D295N **(f)**. Progress curves of c-di-GMP hydrolysis catalyzed by PdeL (0.25 μM) were acquired (See: Materials and methods) with chromatogram peak areas converted to nucleotide concentrations according to calibration. Substrate (c-di-GMP) and product (pGpG) are indicated by downwards and upwards triangles, respectively. Grey circles correspond to their sum. **(e)** Wild-type PdeL exhibits biphasic progress curves. Black lines represent theoretical progress curves obtained by fitting of the model (**Fig. S5a**) to the data. Kinetic parameters are given in **Fig. S5b. (f)** PdeL D295N mutant exhibits linear progress curves that can be fitted with a simple Michaelis-Menten model yielding a k_cat_ of 0.30 s^-1^. **(g)** PdeL catalytic activity is shown as initial velocity at different c-di-GMP concentrations. The activity of PdeL wild type (solid lines) or a 1:1 mix of PdeL wild type and PdeL E141A was determined at the protein concentrations indicated. **(h)** Model for the c-di-GMP mediated control (orange arrows) of transcriptional autoregulation (blue) and catalytic activity (green) of PdeL. Note that c-di-GMP mediated inhibition is not immediate but relies on a slow PdeL R → T transition (see text).

To functionally probe T- and R-states, we aimed to identify mutants that would disfavor one or the other dimer conformation. T- and R-state conformations differ in their dimerization interface, where helices α5, α6, and loop 6 form isologous interactions with the symmetry mates (**Fig. 2c,e, 3b,c**). The strongest difference is seen in the relative positions of helices α5* and symmetry related α6, which are aligned laterally in the R-state but form a head-to-head arrangement in the T-state (**Fig. 3b,c**). The latter arrangement is electrostatically unfavorable but is stabilized by the side-chain of D295, which is capping the N-termini of both helices (**Fig. 3c**). The α5 - loop 6 region structurally couples the dimerization interface to substrate occupancy in the nearby active site. Residue E235 adopts a central hinge function by H-bonding either with active-site residue D263 in the R-state or, upon engagement of D263 in binding of c-di-GMP/Mg^2+^, with residue T270 of the helix α5, which in turn forms H-bonds with Ser298 of the adjacent α6 (**Fig. 3b,c**). In line with this view, disulfide cross-linking of mutants E235A, T279A, and D295N was insensitive to c-di-GMP indicating that the T-state of these mutants is destabilized and the R⟷T equilibrium is tilted towards the R-state (**Fig. 3a**).

In agreement with these *in-vitro* results, strains expressing mutants locked in the R-state showed de-repressed *pdeL* transcription, irrespective of the cellular c-di-GMP concentration (**Fig. 3d**). Moreover, a random genetic screen for *pdeL* alleles able to restore swimming motility at high c-di-GMP concentrations (**Fig. S4a**) identified a range of activating substitutions in PdeL, including T270A and D295N (**Fig. S4b**). Most of these spontaneous mutations are positioned within or close to the dimerization helices α5 and α6 (**Fig. S4c**) and caused de-repression of *pdeL* transcription (**Fig. S4d**), arguing that destabilization of the T-state results in hyperactive PdeL.

Altogether, this demonstrated that PdeL-mediated transcription activity inversely scales with c-di-GMP, while mutants locked in the R-state retain maximal activity even under high c-di-GMP conditions.

### Destabilizing the PdeL T-state conformation results in enhanced catalytic activity

Next, we asked if the observed R⟷T conversion also affects PdeL catalysis. D295, the residue critical for stabilization of the PdeL T-state (**Fig. 3c**), is conserved in a large fraction of bacterial EAL phosphodiesterase domains (**Fig. S4e,f**). This raised the possibility that the R⟷T switch represents a more common allosteric feedback mechanism of EAL proteins and prompted us to test enzyme activity of wild type and mutant forms of PdeL. Phosphodiesterase activity assays showed that the initial velocity of the D295N R-lock mutant was about four times higher than wild type (**Figs. 3e, f**). Notably, while the progress curve of the D295N variant was consistent with simple Michaelis-Menten enzyme kinetics with a low K_m_ and a k_cat_ of 0.30 s^-1^, the wild-type enzyme showed a gradual decrease in velocity towards a linear phase over about 1500 seconds. We established an R-to-T equilibrium-based kinetic model (**Fig. S5a**), which is in good agreement with the data in Fig. 3e. For the fit, k_cat,R_ was restrained to the D295N value of 0.30 s^-1^ and a 10-fold tighter binding of the substrate to the T-state was assumed. Under these conditions, the fit yields a T/R equilibrium constant of 5.6 and a k_cat,T_ of 0.02 s^-1^ **(Fig. S5b)**, indicating that at low enzyme concentrations and in absence of substrate, 15% of the PdeL molecules adopt an R-state conformation, which is 15-times more active than the T-state. The slow response of wild-type PdeL to c-di-GMP is unusual and likely reflects slow R⟷T transitions as suggested by the kinetic model **(Fig. S2c,d)**. Note that slow kinetics was also observed in the MST redistribution experiment (**Fig. 2b**). It is plausible that *in vivo* this behavior delays changes in *pdeL* transcription upon fluctuating levels of c-di-GMP.

The activity of PdeL may be reduced by c-di-GMP due to negative cooperativity between two active sites of one dimer. If so, PdeL heterodimers consisting of a wild-type and mutant protomer unable to bind c-di-GMP should be at least partially protected from this inhibitory effect. To test this, we compared the activity of purified wild-type PdeL with a stoichiometric mix of PdeL wild type and the E141A mutant, which is unable to bind c-di-GMP. While both fractions showed similar activities at low substrate concentrations, heterodimers indeed displayed significantly higher turnover rates at higher substrate concentrations **(Fig. 3g)**. These findings provide an entry point into understanding how c-di-GMP inhibits PdeL activity.

In summary, PdeL transcriptional and catalytic activities inversely scale with c-di-GMP, while mutants locked in the R-state retain maximal transcriptional and enhanced catalytic activity even under high c-di-GMP conditions. Based on this, we propose that at increasing concentrations of c-di-GMP the equilibrium of PdeL is pushed from the R- to the T-state, leading to tetramer dissociation and the concomitant obstruction of transcriptional autoregulation and catalysis (**Fig. 3h)**.

### PdeL imposes binary c-di-GMP signaling regimes with memory

Above we showed that PdeL activity and *pdeL* transcription inversely scale with c-di-GMP (**Fig. 1f**). To address how the *pdeL* promoter responds to dynamic changes of c-di-GMP, we monitored a *pdeL*-*lacZ* reporter in a strain engineered to tune c-di-GMP **(Figs. 1f, 4a)**. When cellular levels of c-di-GMP were gradually increased (by lowering IPTG concentrations), *pdeL* expression remained constantly high and only turned off in the low micromolar range of c-di-GMP **(Fig. 4b)**. Tuning c-di-GMP levels in the opposite direction led to the activation of *pdeL* transcription but exposed a large hysteresis window covering the nano- and sub-micromolar range of c-di-GMP. For example, at 1 µM c-di-GMP, *pdeL* transcription remained ON in cells that had experienced low c-di-GMP before but remained OFF in cells with a history of high c-di-GMP levels **(Fig. 4b)**. History-dependent differences in *pdeL* transcription were largest at sub-micromolar c-di-GMP concentrations, closely reflecting physiological c-di-GMP distributions in wild-type under similar growth conditions (Reinders et al., 2015). As expected, the hysteresis window gradually narrowed over time following c-di-GMP equilibration **(Fig. 4c)** and completely collapsed in mutants lacking the CIB motif **(Fig. 4d)**, reinforcing the idea that binding of PdeL to CIB largely accounts for cooperativity and the switch-like behavior of *pdeL* expression. Hysteresis of *pdeL* expression appears to be mediated by gradual promoter activation when c-di-GMP levels decrease and by a sharp drop of promoter activity followed by protein dilution when c-di-GMP levels are increased **(Figs. 4e,f)**. Direct measurements of c-di-GMP in *E. coli* populations, in which *pdeH* expression is tuned with IPTG in both directions exposed stably divergent c-di-GMP concentrations. These were dependent on PdeL and were only observed 4.5 hours after shifting cells to the new IPTG concentrations **(Fig. 4g)**, while the concentration differences had collapsed eight hours after the shift **(Fig. 4h)**.

**Figure 4:**
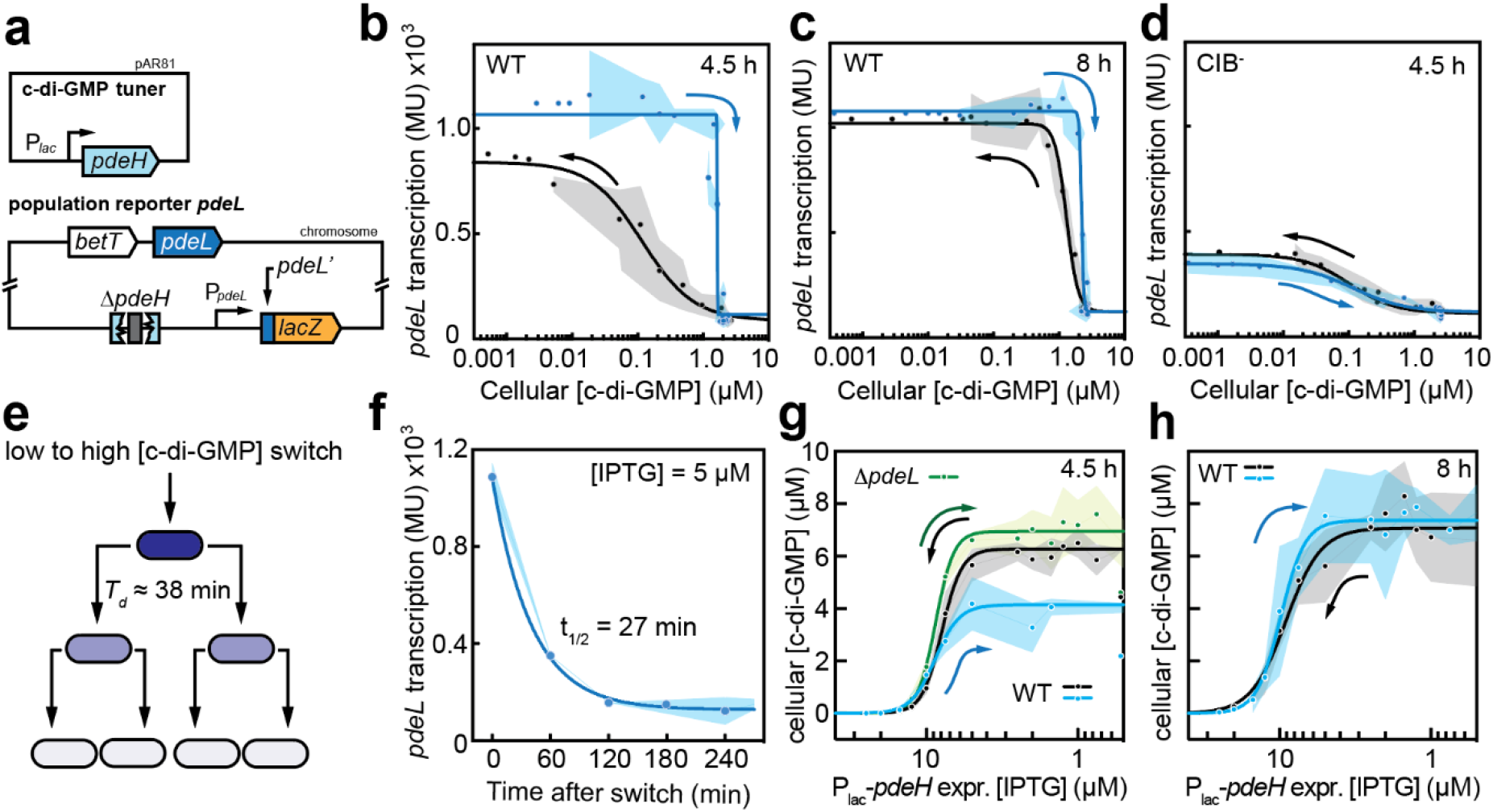
Expression of *pdeL* is bistable. **(a)** Schematic of plasmid and reporter strain used to determine *pdeL* expression at different c-di-GMP concentrations. The *pdeL* promoter region was fused to *lacZ* by replacing the native *lac* promoter in the chromosome (Reinders et al., 2015). To tune levels of c-di-GMP, the chromosomal copy of *pdeH* was deleted and replaced by an IPTG-inducible copy on plasmid pAR81. **(b,c)** Hysteresis experiments with reporter strain shown in (a). LacZ reporter activity was determined 4.5 hours **(b)** and 8 hours **(c)** after equilibrating c-di-GMP levels in the reporter strain. Arrows indicate the direction of changes in c-di-GMP concentration from pre-established low (blue) or high (black) c-di-GMP levels. Curves were fitted with a sigmoidal least square fit. **(d)** Hysteresis experiment with a strain containing a mutated CIB box in the *pdeL* promoter upstream of the *lacZ* reporter. **(e)** Schematic of protein dilution during cell division with the doubling time recorded for the conditions used. **(f)** Time-dependent *pdeL* OFF-kinetics. The reporter strain (a) containing the c-di-GMP tuner plasmid was grown with 65 µM IPTG overnight and diluted back into fresh medium with 5 µM IPTG before monitoring *pdeL* transcription. Data points were fitted with a simple decay model yielding a half-life of t_1/2_ = 27 min. **(g,h)** Development of c-di-GMP concentrations upon altered expression of the c-di-GMP tuner (a) in strains with pre-established low (blue, green) or high (black) c-di-GMP levels. Black and blue lines indicate the behavior of a *pdeL*^*+*^ strain, green marks the *ΔpdeL* mutant. C-di-GMP levels were measured 4.5 hours **(g)** or 8 hours **(h)** after equilibrating the c-di-GMP tuner by adjusting the IPTG concentration.

Thus, both *pdeL* transcription and PdeL-mediated c-di-GMP levels show bistability, indicating that PdeL is a molecular switch that confers cellular memory and robustly determines cellular levels of c-di-GMP in *E. coli*.

### Bimodal *pdeL* expression imposes binary c-di-GMP signaling regimes in individual cells

Next, we asked if the observed bistability of *pdeL* expression establishes bimodal c-di-GMP regimes in individual cells. To assay *pdeL* promoter activity at the single-cell level and at different concentrations of c-di-GMP, we engineered transcriptional fusions to different fluorophores and monitored them in the strain harboring a tunable copy of *pdeH* **(Fig. 5a)**. While the *pdeL*::*gfp* reporter was off in all cells at high c-di-GMP and homogenously active at low c-di-GMP concentrations, bimodal expression was observed at intermediate c-di-GMP levels **(Fig. 5b,c)**. To test if bimodal *pdeL* expression correlated with bimodal patterns of c-di-GMP, we made use of a novel c-di-GMP biosensor that was engineered by fusion of a circularly-permutated GFP (Nadler et al., 2016) to the c-di-GMP-binding domains of BldD from *Streptomyces* (Tschowri et al., 2014). Fluorescence intensity of this sensor directly scales with c-di-GMP levels with maximal and minimal intensities above 1 µM and below 100 nM, respectively. Recording of the GFP-based c-di-GMP sensor and a chromosomal *pdeL*::mCherry reporter in the same strain at intermediate c-di-GMP levels revealed strictly inverse fluorescence patterns indicating that expression of *pdeL* drives bimodal distributions of c-di-GMP in populations of *E. coli* **(Fig. 5d)**. To corroborate the idea that *pdeL* imposes bistability and bimodality on bacterial populations, we compared the dynamic changes of c-di-GMP in individual cells of a *pdeL*^+^ and a Δ*pdeL* strain upon gradually changing c-di-GMP levels using the *P*_*lac*_*-pdeH* tuner plasmid. Strains were grown overnight in the absence of IPTG to pre-establish high intracellular c-di-GMP concentrations and were then shifted to fresh medium containing gradually different IPTG concentrations leading to increased expression of *pdeH*. Increasing IPTG concentrations (decreasing c-di-GMP levels) led to a gradual and monomodal reduction of c-di-GMP levels in the Δ*pdeL* strain. In contrast, an isogenic *pdeL*^*+*^ strain displayed two distinct subpopulations individual cells showing either high or low, but no intermediate c-di-GMP concentrations **(Fig. 5e)**. The PdeL-mediated switch of a subfraction of *E. coli* cells to a low c-di-GMP levels occurred at IPTG concentrations that are significantly below those required to reduce c-di-GMP in the absence of PdeL. Thus, PdeL is a hypersensitive module that converts gradual changes of c-di-GMP into a binary outcome.

**Figure 5:**
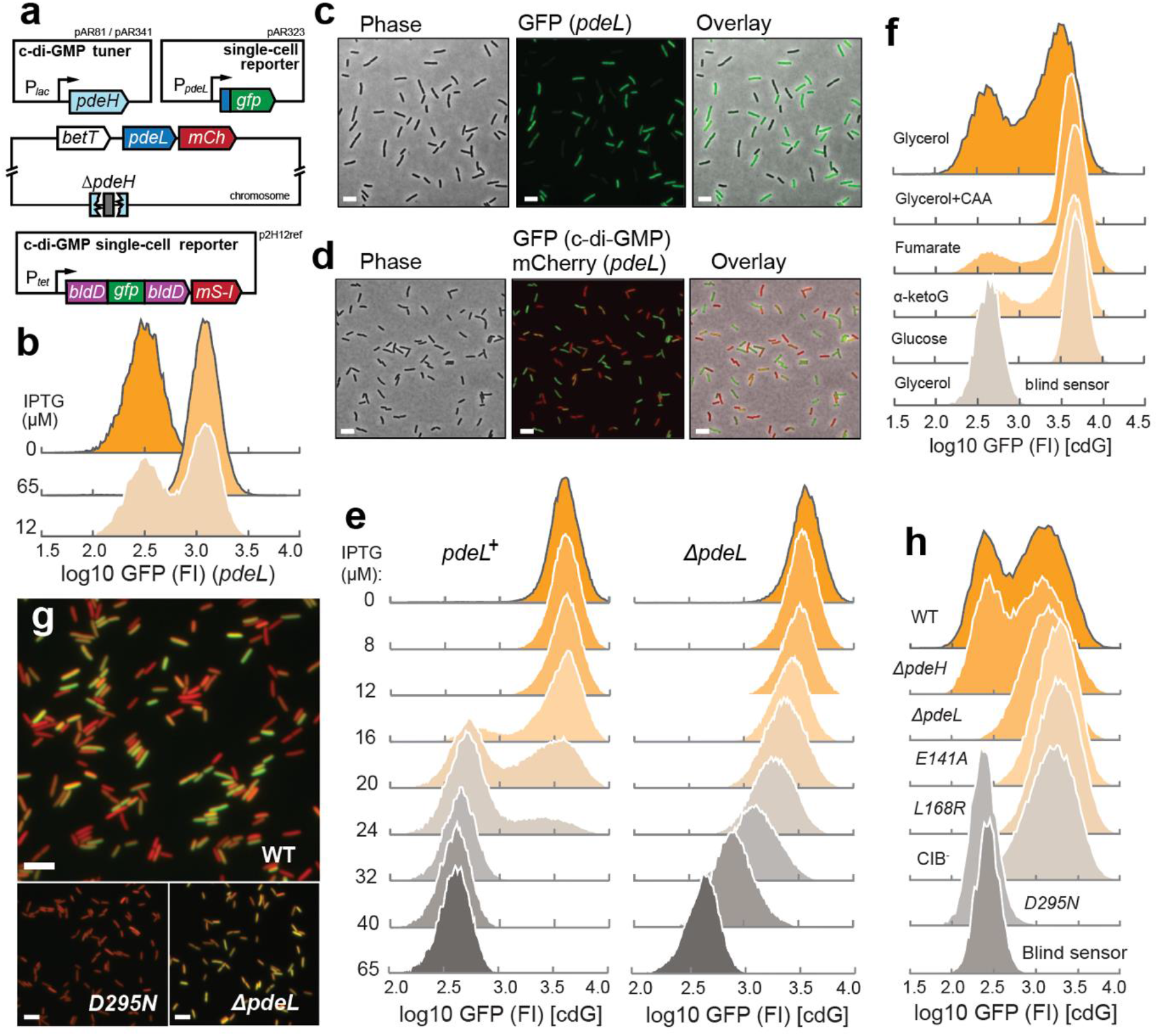
Stochastic expression of *pdeL* established bimodal c-di-GMP regimes. **(a)** Schematic of reporter constructs used. Single-cell *pdeL* transcription was measured with a transcriptional fusion to *mCherry* (*mCh*) downstream of the *pdeL* gene on the chromosome of *E. coli* (CGSC 7740) or with a plasmid-born *P*_*pdeL*_-*gfp* construct (pAR323). To tune c-di-GMP levels, a P_lac_-driven copy of *pdeH* was expressed from plasmid pAR81. Expression of the operon containing the cdG reporter and mScarlet-I (*mS-I*) was induced with 200 nM anhydrotetracycline from plasmid p2H12ref. **(b)** *pdeL* promoter activity was determined in a *pdeL*^+^ Δ*pdeH* mutant of *E. coli* (CGSC 7740) harboring plasmids pAR81 (c-di-GMP tuner) and pAR323 (*pdeL* reporter). Cultures grown in the absence of IPTG (high c-di-GMP) were shifted to media containing 0, 12, or 65 µM IPTG, grown for 6 hrs and analyzed by flow cytometry. **(c)** Fluorescence micrographs illustrating bimodal *pdeL* promoter activity in an *E. coli* strain (CGSC 6300) carrying the GFP reporter plasmid pAR323. Scale bar: 5 µM. **(d)** Fluorescence micrographs illustrating *pdeL* expression (red) and c-di-GMP levels (green) in *E. coli* strain CGSC 6300 carrying a chromosomal *pdeL*::*mCherry* reporter and a plasmid-born copy of the cdG reporter (p2H12ref). Scale bar: 5 µm. **(e)** PdeL establishes binary c-di-GMP regimes. An *E. coli* (CGSC 7740) Δ*pdeH* strain carrying plasmid pAR81 was used to tune c-di-GMP levels in a *pdeL*^+^ and Δ*pdeL* mutant background. Strains also carried plasmid p2H12ref and were precultured under conditions that establish high intracellular c-di-GMP levels (0 µM IPTG). Cultures were then diluted into media containing increasing levels of IPTG, were grown for 8 hrs and analyzed by flow cytometry. **(f)** *E. coli* strain CGSC 7740 containing plasmid p2H12ref was grown in minimal media with different carbon sources to mid log phase and analyzed by flow cytometry. A ‘blind’ sensor carrying a mutation in the c- di-GMP binding site of BldD was used as a control. **(g)** Fluorescence micrographs illustrating c-di-GMP levels in *E. coli* wild type strain CGSC 6300 (WT), *pdeL D295N*, and Δ*pdeL* mutant cells carrying plasmid p2H12ref. Cultures were grown in glycerol minimal medium. Overlay of green: c-di-GMP (GFP), and red: constitutive *mScarlet-I* expressed from the same operon as the c-di-GMP sensor from p2H12ref. Scale bar: 5 µM. **(h)** Distribution of c-di-GMP in *E. coli* populations carrying plasmid p2H12ref and grown in glycerol-based minimal medium. Mutants included Δ*pdeL, pdeL E141A* (substrate binding), *pdeL L186R* (tetramerization), *pdeL D295N*, randomized CIB box (CIB^-^). Sensor control was as indicated in (f).

To scrutinize the role of *pdeL* on c-di-GMP levels under more physiological conditions that do not depend on artificial c-di-GMP tuning, we sought to monitor c-di-GMP levels in individual cells of *E. coli* MG1655 (CGSC 6300) (Guyer et al., 1981). In contrast to its derived genotype CGSC 7740, which has artificially low levels of c-di-GMP due to an IS insertion in the *flhDC* promoter region (Barker et al., 2004; Blattner et al., 1997) leading to the constitutive expression of flagellar genes and of *pdeH* (Reinders et al., 2015), the *flhDC* promoter region is uncompromised in the CGSC 6300 background and its c-di-GMP metabolism not genetically altered (Barker et al., 2004, Kim et al., 2020).

When strain CGSC 6300 was grown on glucose-based minimal media, c-di-GMP levels were uniformly high **(Fig. 5f)**. In contrast, bimodal patterns of c-di-GMP were observed when this strain was grown in minimal medium with alternative carbon sources like glycerol, fumarate, or alpha-ketoglutarate **(Fig. 5f)**. The addition of casamino acids to minimal glycerol media abolished c-di-GMP bimodality, phenocopying the monomodal distribution observed on glucose. Importantly, c-di-GMP bimodality in glycerol media was strictly dependent on PdeL, but not on PdeH **(Figs. 5g,h)**. Mutants abolishing PdeL enzyme activity (E141A) or tetramerization (L168R) as well as mutations of the CIB box showed monomodal high c-di-GMP distributions. These data are in full accordance with the mechanistic and regulatory model proposed above. Accordingly, a mutation that locks PdeL in the active R-state (D295N) also abolished bimodality but generated cells with low c-di-GMP **(Fig. 5g,h)**. Thus, PdeL converts stochastic fluctuations of c-di-GMP into a robust binary output with subpopulations maintaining either low or high levels of c-di-GMP.

### PdeL instructs *E. coli* lifestyle changes and protects against phage predation

Based on the observed c-di-GMP bimodality, we set out to probe if PdeL impacts the behavior of cells experiencing changes in c-di-GMP. For this, we assayed *E. coli* biofilm formation and escape, processes that are directly controlled by c-di-GMP (Hengge, 2021; Jenal et al., 2017; Steiner et al., 2013). First, we monitored the effect of PdeL on poly-GlcNAc-dependent (PGA) biofilm formation (Steiner et al., 2013) in a strain harboring the IPTG tunable *pdeH* construct. When c-di-GMP levels were gradually increased by lowering *pdeH* expression, surface attachment was triggered only when IPTG concentrations reached levels at which *pdeL* expression is turned off. In contrast, an isogenic strain lacking PdeL showed biofilm formation already at higher *pdeH* expression levels **(Fig. 6a)**. Thus, PdeL effectively buffers against c-di-GMP noise to sustain the planktonic lifestyle of *E. coli*. Conversely, PdeL strongly promoted the escape of *E. coli* from pre-established biofilms upon lowering c-di-GMP levels. The observed large variations of escape rates are in line with a highly stochastic induction of PdeL under these conditions **(Fig. 6b)**.

**Figure 6:**
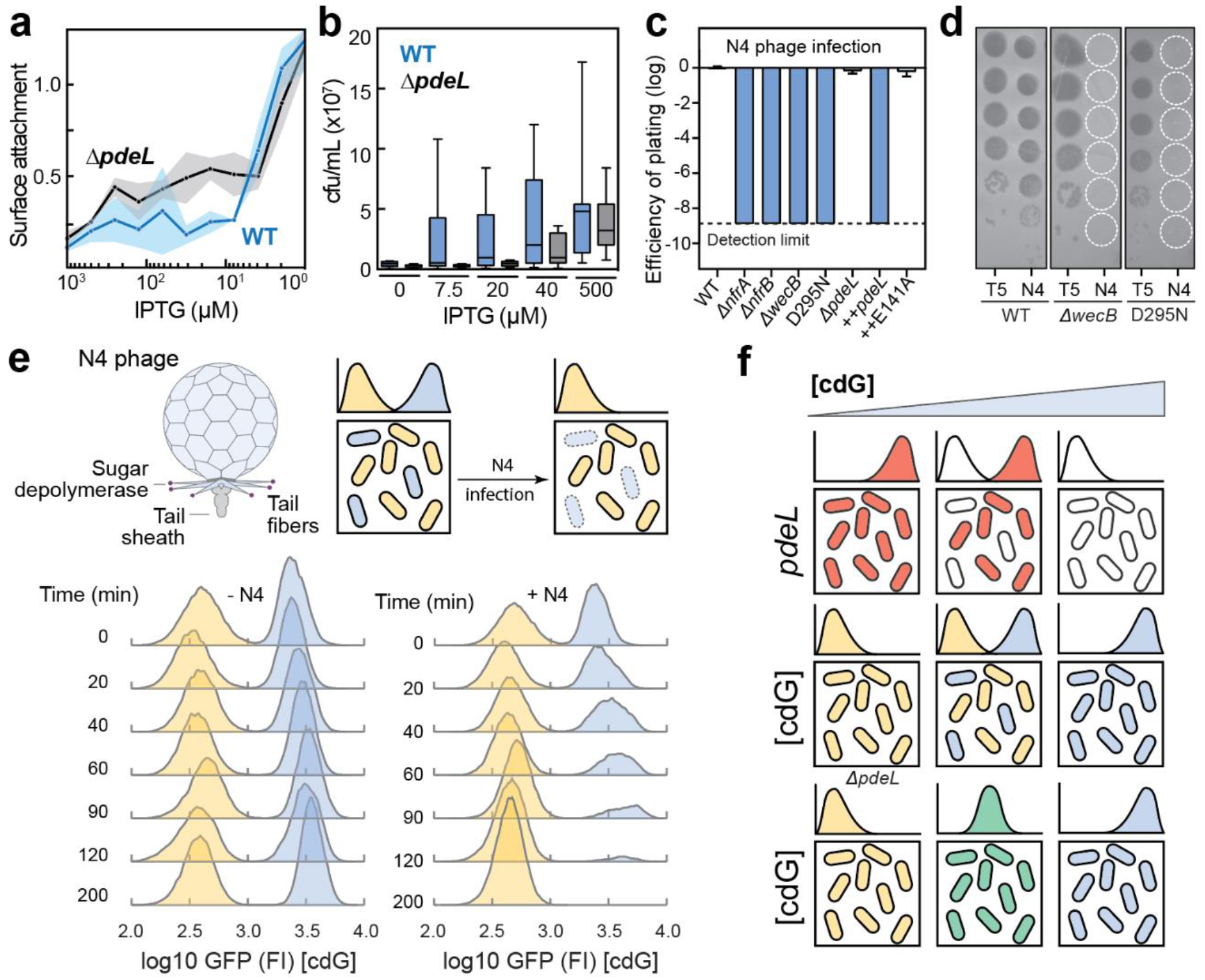
PdeL controls c-di-GMP-dependent biofilm formation and escape and protects against phage N4. **(a)** Biofilm formation of *E. coli* wild type (blue) and *ΔpdeL* mutant (black) at different intracellular levels of c-di-GMP. The concentration of c-di-GMP was set in the biofilm assay strain with plasmid pAR81 (see: Fig. 5a and Materials and Methods) by IPTG induction. **(b)** Escape of *E. coli* wild type (blue) and *ΔpdeL* mutant (black) cells from pre-formed biofilm upon IPTG-mediated induction *pdeH* from the c-di-GMP tuner plasmid pAR81. The number of cells escaping to the planktonic fraction was scored 3 hrs after fresh medium with different concentrations of IPTG was added to pre-formed biofilms. Median values are indicated with error-bars showing upper and lower quartiles. **(c)** PdeL protects against infection by phage N4. EOP (efficiency of plating) depicts the phage susceptibility of mutants relative to *E. coli* wild-type strain CGSC 6300. Strains with deletions in known N4 infection genes (*nfrA, nfrB* and *wecB*) or with mutations in *pdeL* are as indicated. Strains expressing *pdeL* (++pdeL) or *pdeL E141A* (++E141A) from an IPTG inducible promoter on plasmid pNDM220 are marked (n = 3). **(d)**. Representative images of plaques formed by phages T5 or N4 on *E. coli* wild-type strain CGSC 6300 with 10-fold serial dilution top to bottom. **(e)** PdeL protects from N4 infection. Top: Schematic drawing of bacteriophage N4 and of the experimental setup. Bottom: *E. coli* wild-type (CGSC 6300) and a *pdeL D295N* mutant harboring the c-di-GMP sensor plasmid were mixed 1:1 and analyzed over time with (right panel) and without phage N4 (left panel). C-di-GMP levels are indicated on the x-axis as fluorescence of the c-di-GMP sensor. **(g)** Model for PdeL control and its impact on cellular c-di-GMP concentrations. Transcription of *pdeL* is activated when c-di-GMP levels drop below a threshold (red cells) and is stalled at increased c-di-GMP levels (white cells). Cells with active pdeL transcription show increased PdeL concentration, which in turn lowers the cellular levels of c-di-GMP (yellow). In contrast, c-di-GMP increase in cells with stalled *pdeL* transcription (blue). Noise-induced fluctuations of c-di-GMP levels result in stochastic expression of *pdeL* establishing bimodal populations harboring distinct concentrations of c-di-GMP. Strains lacking PdeL exclusively adopt unimodal distributions of c-di-GMP levels (green).

Next, we reasoned that c-di-GMP-mediated production of surface exopolysaccharides may serve as an entry door for bacteriophages and that stochastic activation of PdeL may serve to protect *E. coli* from phage killing. Specific members of these important bacterial predators use exopolysaccharides, capsules, or LPS as primary receptors and have evolved virion-associated polysaccharide depolymerases to gain access to secondary receptors on the cell surface (Nobrega et al., 2018). Screening a library of *E. coli* bacteriophages (Maffei et al., 2021) for agents that specifically prey on cells with high levels of c-di-GMP led to the identification of the podovirus N4 **(Fig. 6e)**, which readily infects *E. coli* strain MG1655 (CGSC 6300) but does not kill an isogenic strain that constitutively expresses *pdeL* **(Fig. 6c)**. Importantly, replacing the *pdeL* wild-type allele on the chromosome with *pdeL D295N* encoding a PdeL R-lock variant lead to complete resistance against phage N4, similar to deletions of known N4 resistance genes like *nfrA, nfrB*, or *wecB* (Kiino and Rothman-Denes, 1989; Kiino et al., 1993; Mutalik et al., 2020) **(Figs. 6c,d)**. The observation that constitutive expression of *pdeL* wild type, but not *pdeL E141A* encoding a catalytically inert variant, conferred phage resistance indicated that PdeL phosphodiesterase activity is critical for phage protection and that N4 specifically infects *E. coli* cells that harbor high levels of c-di-GMP. This was confirmed when cells of *E. coli* wild type and the *pdeL D295N* mutant were mixed and challenged with N4 phages. As shown in **Fig. 6e**, N4 selectively infected cells with inactive PdeL and high levels of c-di-GMP. Together, these experiments emphasize the relevance of PdeL as a central regulatory component of the motile-sessile switch in *E. coli* and suggest that stochastic activation of PdeL effectively protects *E. coli* from phage predation.

## Discussion

### PdeL is a binary switch digitalizing cellular c-di-GMP levels

Binary switches generally operate on the transcriptional level to modulate the cell’s genetic circuitry (Carraro et al., 2020). Here, we describe a molecular switch that converts gradual changes of a small and highly potent signaling molecule into robust binary signaling modes (**Fig. 6f**). We propose that by digitalizing c-di-GMP, *E. coli* elegantly evades ambiguous signaling states that can be caused by noise or by converging input from multiple players of complex networks and that could otherwise result in intermediate, non-productive levels of the second messenger. Establishing stable binary states may be particularly important for all small nucleotide-based signaling molecules like cAMP or c-di-GMP because they are often part of complex regulatory networks and because they regulate many cellular processes at the post-translational level and on very short time scales. Tight binary control of potent, high affinity NSMs might be of general importance for bacteria to sustain short response times and by that maintain their functional versatility and overall fitness. For instance, when encountering surfaces, planktonic bacteria respond to mechanical cues by increasing their c-di-GMP or cAMP levels. This blocks motility (Boehm et al., 2010; Schniederberend et al., 2019), leads to the assembly of adhesion factors (Hug et al., 2017; Laventie et al., 2019) and to the expression of virulence genes (Persat et al., 2015). Molecular switches like PdeL could define an upper threshold of c-di-GMP that maintains the ability of planktonic bacteria to respond to mechanical cues in a highly sensitive manner when encountering surfaces.

In addition to keeping a tight grip on the global signaling program, switches like PdeL may also contribute to more specific signaling by effectively insulating local signaling nodes (Hengge, 2021). The observation that mutants lacking individual diguanylate cyclases or phosphodiesterases can have highly specific phenotypes without manifest changes in global c-di-GMP together with the finding that such components physically interact with their c-di-GMP targets, had suggested that bacteria are able to spatially sequester individual signaling systems and use c-di-GMP as local pacemaker to regulate cellular processes in a highly specific manner (Collins et al., 2020; Lori et al., 2015; Richter et al., 2020; Ross et al., 1987; Sarenko et al., 2017). To effectively insulate local signaling modules, the global c-di-GMP pool needs to be kept low by a master phosphodiesterase. Such a role is for instance played by PdeH, which is co-expressed with flagellar genes in *E. coli* and strongly reduces the global pool of c-di-GMP under conditions favoring motility (Barker et al., 2004; Reinders et al., 2015). Similarly, stochastic regulation of PdeL could switch *E. coli* between a state of low overall c-di-GMP and high local signaling specificity and a state of globally high c-di-GMP where local modules would be overruled by the global cellular program. If so, PdeL would elegantly couple the transition between local and global signaling to the prevailing c-di-GMP concentration and by that allow *E. coli* to effectively veto local signaling independence under conditions where all cellular processes need to be in line with a high c-di-GMP program.

### Molecular digitizer PdeL imposes population heterogeneity

Binary signaling modes generated by a hypersensitive switch naturally impose heterogeneity in populations experiencing dynamic environmental changes and behavioral transitions. This may serve as bet-hedging strategy to protect part of the population from predators like phage N4 that exploit conserved surface structures associated with certain behavior or physiological states. Given the prominent role of c-di-GMP in stimulating EPS synthesis and secretion (Krasteva et al., 2010; Merighi et al., 2007; Steiner et al., 2013; Thongsomboon et al., 2018; Whitney et al., 2012), high c-di-GMP states may impose a specific predator-mediated burden and predispose bacteria for phage infections. In line with this, we have recently shown that N4 uses a novel surface glycan in *E. coli* as primary receptor that we termed NGR (N4 glycan receptor). The production of NGR strictly depends on c-di-GMP, providing an explanation for the strong protective role of PdeL against N4 infections (Sellner et al., 2021). Stochastic modulation of surface structures and population heterogeneity may help *E. coli* and other bacterial pathogens to circumvent host immunity. For instance, c-di-GMP was shown to modulate O-antigen distribution of the most outer layer of lipopolysaccharide in *P. aeruginosa* and by that contributes to immune evasion of this human pathogen (McCarthy et al., 2017). Similarly, stochastic control and heterogeneity of metabolic rates or virulence gene expression was shown to promote resilience or cooperation of bacterial pathogens and by that increase their overall fitness in the host (Brauner et al., 2016; Davis and Isberg, 2018)

In addition to its role in stress mitigation, population heterogeneity may also increase the overall fitness of bacterial populations by generating multiple cellular identities. For instance, during adaptation to surfaces it may be an advantage for planktonic bacteria to functionally differentiate into adherent and non-adherent subpopulations to facilitate surface colonization or to minimize the risk involved in such fundamental lifestyle changes. It was recently proposed, that the opportunistic human pathogen *P. aeruginosa* facilitates adherence and spreading on abiotic and biotic surfaces by generating distinct cell types through asymmetric divisions (Laventie et al., 2019). Although in this particular case distinct cell fates result from an inherent developmental program, generating different cellular identities through stochastic processes likely provides similar fitness advantages. Finally, established biofilm communities may also generate and benefit from heterogenous signaling states when experiencing environmental or nutritional challenges that alter their global c-di-GMP levels. *E*.*g*., to escape from biofilms, bacteria need to reduce their overall c-di-GMP concentration through the activation of one or several phosphodiesterases (Rumbaugh and Sauer, 2020). A hypersensitive stochastic switch like PdeL may amplify such regulatory input and convert it into robust low c-di-GMP states to facilitate the escape of individual members of surface attached communities without compromising the overall integrity of the biofilm.

### Mechanistic basis of the PdeL switch

PdeL is a c-di-GMP-dependent sensor, catalyst and transcription factor at the same time. PdeL activity is highest at low levels of c-di-GMP but is reduced when c-di-GMP levels increase. Our data suggest that the regulatory behavior of PdeL relates to its dynamic transition between inert T-state dimers and active R-state tetramers. The dynamic equilibrium between these two conformations is inversely influenced by the PdeL protein concentration and by c-di-GMP and likely explains the cooperative behavior of PdeL as a transcription factor. The T-to-R-state switch is mediated through tight structural coupling between the substrate-binding site and the dimerization interface of PdeL (Sundriyal et al., 2014). Similar structural coupling was proposed for the activation of phosphodiesterases by associated sensory domains (Rao et al., 2009; Winkler et al., 2014) and for EAL domains that have lost their catalytic function and have adopted a role as c-di-GMP effectors (Minasov et al., 2009; Navarro et al., 2011). This and the observation that residues involved in the conformational switch of PdeL are widely conserved in EAL domains, argues that tight coupling between substrate binding and dimerization could be a general mechanism through which phosphodiesterases are inhibited at high substrate concentrations.

Based on our data we propose that PdeL activates its own transcription by directly recruiting RNA polymerase and possibly via the formation of a tetramer that engages both CIB and CDB upstream of the *pdeL* promoter. Binding to CDB and CIB may increase PdeL concentration locally, thereby shifting the equilibrium towards a highly active tetramer. This may lead to DNA bending and displacement of the general gene silencer H-NS. In line with this, binding sites for IHF and Fis, proteins known to promote DNA bending, are positioned upstream of the *pdeL* promoter between CIB and CDB (Grainger et al., 2006). How exactly PdeL interferes with H-NS and how it is able to recruit RNA Pol to the *pdeL* promoter region remains to be shown.

### Molecular digitizer PdeL opens up new research questions

This study raises several important future questions. How widespread are mechanisms that impose cellular heterogeneity to NSM-based networks? Are molecular mechanisms converting graded changes of NSMs into binary outputs a necessary consequence of or an opportunity of complex signaling architectures of networks operating with small signaling molecules like cAMP, c-di-GMP and others? Also, what is the exact role of PdeL and similar digital converters in spatiotemporal control of NSM networks? *E*.*g*., does the PdeL transcription factor exclusively control its own expression, or does this regulon expand to additional *E. coli* genes? Finally, why is *pdeL* autoregulation strictly coupled to the activity of Cra, a sensor of the metabolic flux through the central carbon metabolism (Folly et al., 2018; Kochanowski et al., 2013)? Cra activity is highest under gluconeogenic conditions and at low growth rates, indicating that c-di-GMP heterogeneity may contribute to population fitness primarily under such growth conditions. The observations that both Cra and c-di-GMP play important roles in regulating virulence of pathogenic *E. coli* (Branchu et al., 2013; Hu et al., 2013; Njoroge et al., 2013; Richter et al., 2014) indicates that important behavioral processes are coordinated with growth and metabolic activity in this organism. How this contributes to the successful adaptation of pathogenic *E. coli* to the human host remains to be determined.

## Materials & Methods

### Bacterial strains and growth media

The bacterial strains and plasmids used in this study are listed in Tables S2 and S3, respectively. *E. coli* K-12 MG1655 (Blattner et al., 1997; Guyer et al., 1981) and its derivatives were grown as indicated in the dedicated methods sections. *E. coli* K-12 MG1655 (CGSC 6300) was obtained from the Coli Genetic Stock Center (CGSC). For strain construction and pre-cultures, LB (Luria Bertani) medium was used. Physiology experiments were performed either in TB (Tryptone Broth; 10 g/l tryptone, 5 g/l NaCl) or M9 minimal medium (Gerosa et al., 2013) to which carbon sources were added from concentrated stock solutions. P1 phage lysate preparation and transduction were carried out as described previously (Boehm et al., 2010).

### Gene deletions and λ**-RED-mediated recombineering**

#### Chromosomal gene deletions and modifications

Gene deletions were essentially carried out either as described (Datsenko and Wanner, 2000) or with the use of a comprehensive mutant library (Baba et al., 2006) and P1 mediated transduction. Chromosomal 3xflag-tagging of genes was carried out according to the published method (Uzzau et al., 2001). For unmodified strains, AB330 (Table S2) or pKD46 was used. pKD46-mediated recombineering was used for construction of strains already harboring chromosomal modifications. Selection markers were removed by site-specific recombination using pCP20 (Datsenko and Wanner, 2000).

#### Construction of promoter-lacZ fusions

Construction of chromosomal promoter-*lacZ* fusions were carried out via λ-RED-mediated recombination as described above. AB989 (Table S2) was used as a recipient strain. The donor PCR fragment harboring the promoter of interest was designed to site-specifically excide P_rha_-*ccdB* and integrate upstream of the native *lacZ* ORF to generate a merodiploid translational fusion. Successful integration events were selected through growth on rhamnose minimal plates.

### Electrophoretic mobility shift assay (EMSA)

5’ Cy3-labeled input DNA was generated either via oligonucleotide annealing or PCR. For oligonucleotides used see Table S4. 10 nM of the input DNA and purified proteins were incubated for 10 min at room temperature in buffer consisting of 50 mM Tris-HCl pH 8.0, 50 mM NaCl, 10 mM MgCl_2_, 10 % Glycerol, 1 mM DTT, 0.01 % Triton X-100, 0.1 mg/mL BSA and 25 µg/mL λ-DNA. As indicated in the Figures, samples were incubated in the presence or absence of 2 mM CaCl_2_ and 50 µM c-di-GMP. Samples were run on 8 % polyacrylamide gel. DNA-protein complexes were analyzed using Typhoon FLA 7000 (GE Healthcare).

### β-galactosidase reporter assay

Strains were grown in TB medium o/n at 37°C. The next day cultures were diluted back 1:500 into fresh medium and grown at 37°C until desired OD_600_. 500 µL of the culture were mixed with 380 µL Z-buffer (75 mM Na_2_HPO_4_, 40 mM NaH_2_PO_4_, 1 mM KCl, 1 mM MgSO_4_) supplemented with 100 µL 0.1 % SDS and 20 µL chloroform. Samples were vortexed for 10 sec and left on the bench for 15 min. 200 µL sample were transferred into a clear 96-well plate. As substrate 25 µL 4 mg/mL 2-nitrophenyl-β-D-galactopyranoside (*o*-NPG) solution (dissolved in Z-buffer) were added. The initial velocity of the color reaction was determined at a wavelength of 420 nm.

### Protein purification

PdeL_EAL_ variants were purified by single StrepII-tag or His-tag affinity purifications, whereas for full-length PdeL and other transcription factors, a heparin purification step was added.

#### StrepII-tag purification

All proteins were cloned into a pET28a vector (Novagen) between NcoI and NotI restriction sites. Proteins were overexpressed in BL21 (AI) cells grown at 30°C in 2 L LB medium. For overexpression of mutant protein variants, the corresponding wild-type version of the gene was deleted in the overexpression strain. At an OD_600_ of 0.6 the culture was induced with 0.1 % L-arabinose. Cells were harvested 4 h post-induction by centrifugation at 6000 g for 30 min at 4°C. The cell pellet was resuspended in 7 mL Buffer A (100 mM Tris-HCl pH 8.0, 250 mM NaCl, 5 mM MgCl_2_, 0.5 mM EDTA, 1 mM DTT) including a tablet of c0mplete mini EDTA-free protease inhibitor and a spatula tip of DNaseI. Cells were lysed by 4 passages of French press, and the lysate cleared at 4°C in a table-top centrifuge set at full speed for 40 min. The cleared supernatant was loaded on 1 mL StrepTactin Superflow Plus resin. The supernatant was reloaded another two times before washing with a total of 50 mL Buffer A. The column was washed with 10 mL Buffer B (100 mM Tris-HCl pH 8.0, 50 mM NaCl, 5 mM MgCl_2_, 0.5 mM EDTA, 1 mM DTT). 500 µL aliquots of proteins were eluted with Buffer B supplemented with 2.5 mM d-Desthiobiotin.

#### His-tag purification

His-tagged proteins, such as Cra-6xHis were expressed at 18°C for 12h with 0.5 mM IPTG. Cells were lysed with 3 passages on the French press, cleared at 35’000g with ultracentrifugation and applied to a 5 ml Ni-chelating column equilibrated with 50 mM Tris pH 8, 1M NaCl, 5mM MgCl_2_. The protein was quantified by recording absorbance at 280 nm and stored at −20 °C at ∼9 mg/ml concentration.

RNA polymerase holoenzyme purification was adapted from (Fong et al., 2010). In brief ELP- intein-α tagged fusion protein and the other core RNAP subunits were co-expressed in terrific broth at an OD_600_ of 0.5-0.7 for 4 h at 37°C via 1mM IPTG and then shifted to 18°C for 12-18 h. Cells were re-suspended in 1.7 ml of pH 8.5 lysis buffer (10 m*M* Tris-HCl [pH 8.5], 2 mM ethylenediaminetetraacetic acid [EDTA], 0.1 mg/mL lysozyme) and lysed by 2 passages of French pressing. The lysate was then cleared by ultracentrifugation at 35’000 x g for 30 min at 4° C. The remaining protocol was performed identically as described in (Fong et al., 2010). *Heparin Purification:* A 1 mL HiTrap Heparin HP was washed with 10 mL H_2_O_dest._, followed by equilibration with 10 mL Buffer B. The eluate from the StrepII-tag affinity purification was loaded three times. After loading, the column was washed with 10 mL Buffer A followed by a washing step with 10 mL Buffer C (100 mM Tris-HCl pH 8.0, 350 mM NaCl, 5 mM MgCl_2_, 0.5 mM EDTA, 1 mM DTT). The protein was eluted in 500 µL fractions with a total of 10 mL Buffer D (100 mM Tris-HCl pH 8.0, 2 M NaCl, 5 mM MgCl_2_, 0.5 mM EDTA, 1 mM DTT). The fractions containing the highest protein concentration were pooled and dialyzed o/n at 4°C against 1.5 L Dialysis Buffer (100 mM Tris-HCl pH 8.0, 250 mM NaCl, 5 mM MgCl_2_, 0.5 mM EDTA, 1 mM DTT). PdeL_EAL_ variants used for cysteine crosslink assays were dialyzed against CXA Buffer (100 mM Tris-HCl pH 7.2, 250 mM NaCl, 2 mM EDTA). The final protein concentration was recorded at 280 nm, and the content of co-purified nucleotide contaminants determined as a ratio of 260/280 nm.

### Microscale thermophoresis

Experiments were carried out with a Monolith NT.115 device using Premium capillaries (both from NanoTemper Technologies). Experiments were performed in PBS supplemented with 5mM DTT, and 0.05% Tween 20 in Premium treated capillaries with 40% LED power and 80% IR-Laser power at 22°C. Laser on and off times was set to 30 and 5 seconds, respectively. In all assays, PdeL-His was labeled using the Monolith NT His-Tag Labeling Kit RED-tris-NTA. Labeled PdeL-His was kept at a constant final concentration of 50 nM. For binding assays, 16 two-fold serial dilutions of an unlabeled partner were used.

To determine the K_d_ of PdeL tetramerization, 1.5-fold serial dilutions of unlabeled PdeL-Strep were prepared starting from 0.21 µM and doped with chromophore labeled PdeL-His (PdeL*) for (time-dependent) MST measurements. The series of PdeL samples was allowed to equilibrate for 2h 30min before adding PdeL* to a final concentration of 50 nM (Fig 3S top panel). Subsequently, the MST profile was measured repeatedly up to 3h 30 min after PdeL* addition. The kinetics of tetramer dissociation was probed by sample dilution (Fig. S2b). For this, an equilibrated series of PdeL samples doped with 50 nM PdeL* were mixed and equilibrated for 4 hrs as above nad then diluted 3-fold with 50 nM PdeL*. K_d_ values were calculated using MO Affinity Analysis software (NanoTemper). Data points were fitted with ProFit 7 (QuantumSoft, Uetikon am See, Switzerland) according to the dimer - tetramer model and an allosteric sigmoidal curve (Y=V_max_*X^h^/(K_half_h + X^h^)) using Prism Graphpad.

### SEC-MALS analysis

PdeL samples (100 µl) varying from 11 to 371 µM were loaded onto a Superdex 200 (10/300) column (GE Healthcare) at constant flow (0.5 ml/min) in 20 mM Tris pH 8.0, 200 mM NaCl, 5 mM MgCl2, 5 mM DTT. The SEC instrument was coupled to an in- line multi-angle light-scattering and differential refractive index detectors (Wyatt Heleos 8+ and Optilab rEX) to measure the apparent mass values (MW_app_) during elution. The inter-detector delay volumes and band broadening, the light-scattering detector normalization, and the instrumental calibration coefficient were calibrated using a standard 2 mg/ml of BSA solution (Thermo Pierce) run in the same buffer, on the same day, according to standard Wyatt protocols. SEC-MALS experiments with PdeL in the presence of the CIB or CDB DNA fragments or Cra protein were performed with the same column and device. In the loaded samples, PdeL concentration was at 94 μM, Cra 94 μM, the DNA fragments CIB and CDB at 120 μM. SEC experiments were performed with the same column using 120 µM PdeL and 240 µM c-di-GMP in the presence of 5 mM CaCl_2_.

### Crystallization

PdeL protein and CIB DNA fragments were mixed with a 2:1 stoichiometry to a final concentration of 100 µM. Crystals were obtained using the sitting-drop vapor-diffusion method after optimizing F7 condition in clear strategy screen (molecular dimension). Crystals were flash-frozen in liquid nitrogen using 20% glycerol as cryoprotectant.

### X-ray diffraction data collection, molecular replacement, and refinement

X-ray diffraction datasets were collected at the Swiss Light Source (SLS) synchrotron on the PXI beamline. Diffraction datasets were processed using XDS (Kabsch, 2010) and the resulting intensities were scaled using SCALA from the CCP4/CCP4i2 suite (Potterton et al., 2018) (Collaborative Computational Project, Number 4, 1994). The structure was solved by molecular replacement (program Phaser (McCoy et al., 2007)) without subsequent refinement using the coordinates of a PdeL-EAL (PDB: 4LYK) monomer as search model. Structural figures were prepared using Dino (http://dino3d.org).

### Mass photometry

The Refeyn OneMP mass photometer was used to determine PdeL oligomerization state at low protein concentrations. 18 μl buffer (20 mM Tris pH 8.0, 200 mM NaCl, 5 mM MgCl_2_) were pre-loaded into a silicone well and mixed with 2 μl of protein (final concentration of 25 nM) prior to data acquisition. 6000 frames were collected using default instrument parameters. DiscoverMP software provided by Refeyn was used for data analysis using default parameters for event extraction and fitting.

### Immunoblotting

Cells were grown in TB medium at 37°C until desired OD_600_. An equivalent of 1 mL of an OD_600_ of 1.0 was pelleted and resuspended in 100 µL SDS Laemmli buffer. Cells were lysed by boiling the sample at 98°C for 10 min. 8 µL of total cell extracts were loaded onto a 12.5 % SDS-polyacrylamide gel, and proteins transferred using a wet blot system. Proteins with 3xFlag-tag were detected with a 1:10.000 dilution of monoclonal mouse α-Flag monoclonal antibody and a 1:10.000 dilution of polyclonal rabbit α-mouse horseradish-peroxidase (HRP) secondary antibody. Proteins were visualized with enhanced chemiluminescence (ECL) detection reagent and imaged in a gel imager (GE ImageQuant LAS 4000).

### Phosphodiesterase enzyme assay

#### Phosphate Sensor assay

Phosphate sensor assay was essentially performed as described in (Reinders et al., 2015). Conversion of c-di-GMP into pGpG was measured indirectly by a coupled alkaline phosphatase (AP)/phosphate sensor online assay. The terminal phosphate of the pGpG product is cleaved by the coupling enzyme AP (20 U/µl, Roche), and the phosphate concentration is determined from the fluorescence increase through binding of phosphate to the phosphate sensor (0.5 µM; Thermo Fisher). PdeL and c- di-GMP concentrations were used, as shown in the individual experiments. Fluorescence increase was detected by excitation at 430 nm and emission at 468 nm. *FPLC assay*: Assays were performed as described in (Sundriyal et al., 2014). Enzymatic activity was assayed offline by FPLC-based steady-state nucleotide quantification following incubation for varying durations. Enzymatic reactions were carried out at 20°C in 100 mM Tris-HCl pH 8.0, 250 mM NaCl, 5 mM MgCl2, 0.5 mM EDTA, 1 mM DTT and 50 M thiamine pyrophosphate as FPLC standard. PdeL and c-di-GMP concentrations were used as described in the result section. The reaction was started by addition of enzyme to a total reaction volume of 600 µl. Samples volumes of 100 µl were withdrawn and the reaction was stopped at different time points by addition of 10 µl of 100 mM CaCl2 and subsequent heating at 98°C for 10 min.

The samples were then analyzed using ion-exchange chromatography (1-mL Resource-Q column) after addition of 890 µL 5 mM ammonium bicarbonate (NH_4_CO_3_) to increase the volume to 1 mL. 500 µL of this was then loaded onto the column. The column was washed thoroughly and the bound nucleotides were eluted with a linear NH_4_CO_3_ gradient (5 mM to 1 M) over 17 column volumes. The amount of pGpG product was determined by integration of the corresponding absorption (253 nm) peak after normalization of the data with respect to the internal thiamine pyrophosphate standard.

#### Online ion-exchange chromatography (oIEC) assay

to acquire automatically quantitative PdeL catalyzed enzyme progress curves by repetitive cycles of sample loading followed by salt gradient elution (0 - 1 M (NH4)_2_SO_4_). The reaction was started by addition of c-di-GMP (final concentration 100 μM) to 0.25 μM PdeL in reaction buffer (100 mM Tris·HCl, pH 8.0, 250 mM NaCl, 5 mM MgCl_2_, 0.5 mM EDTA, 1 mM DTT). This was followed by sequential aspiration of reaction mix aliquots to the column at defined time points followed by a salt gradient elution (0 - 1 M (NH4)_2_SO_4_). Note that when the sample arrives on the column the reaction is stopped without the need of further intervention due to substrate immobilization. The chromatogram peaks corresponding to the nucleotides were fitted by Gaussian distributions using a custom made automized routine implemented in ProFit 7 (QuantumSoft, Uetikon am See, Switzerland). Peak areas (mAU. ml) were converted to molar nucleotide amounts using a scale factor obtained by calibration with a set of serially diluted c-di-GMP samples. Progress curves were fitted using numeric integration of the differential equations corresponding to the kinetic model shown in Fig. S5a implemented as a Python script in ProFit 7.

### Cysteine crosslink assay

10 µM PdeL_EAL_-3xFlag-StrepII variants purified in CXA Buffer (100 mM Tris-HCl pH 7.2, 250 mM NaCl, 2 mM EDTA) were incubated for 10 min at room temperature in the presence of 10 mM CaCl_2_, either in presence or absence of 50 µM c-di-GMP. Proteins were crosslinked for 1 h at room temperature with an 8-fold molar excess (80 µM) of bismaleimidoethane (BMOE). Crosslink reaction was quenched for 15 min at room temperature by addition of 50 mM DTT. Samples were supplemented with SDS Laemmli buffer and proteins denatured by heating at 98°C for 5 min. Samples were loaded on a 1.5 mm thick 12.5 % SDS-polyacrylamide gel and detected by staining with Coomassie Brilliant Blue G according to the staining protocol from Sigma (product number B2025). In brief: Gel was fixed for 30 min in Fixing Solution and stained over night at room-temperature in staining solution. Gel was destained with 10 % acetic acid (v/v), 25 % (v/v) methanol for 60 sec with shaking. Gel was rinsed with 25 % methanol and destained for 24 h with fresh 25 % methanol.

### Microscopy

Cells were placed on a PBS pad solidified with 1% agarose. Fluorescence and differential interference contrast (DIC) microscopy were performed on a DeltaVision Core (Applied Prescision, USA) microscope equipped with an Olympus 100X/1.30 Oil objective and an EDGE/sCMOS CCD camera. Exposure time for microscopy picture was 0.05 sec for bright field (POL), 0.1 sec for GFP for the c-di-GMP sensor, 0.2 sec for *pdeL-gfp* and 0.4 or 0.5 sec for *pdeL-mCherry*. For all settings, the ND filter was set to 100% transmission. For strains grown on M9 glycerol, microscopy was performed using an inverted microscope (Eclipse Ti2, Nikon Instruments Europe B.V.) with 100x oil immersion objective (CFI Plan Apo λDM100x Oil, Nikon). Exposure time was set to 0.05 sec for phase contrast, 0.2 sec for GFP, and 0.2 sec for mCherry.

### C-di-GMP measurements

C-di-GMP measurements were performed according to the published procedure (Spangler et al., 2010). *E. coli* cells were grown in 24 mL TB medium at 37°C to an OD_600_ of 0.5. Cells were pelleted and washed in 300 µL ice-cold distilled water. After washing, the cell pellet was resuspended in 300 µL ice-cold extraction solvent (acetonitrile/methanol/H_2_O_dest._, 40/20/20 v/v/v). After pelleting, the supernatant was transferred to a 2 mL safe-lock tube, and the extraction procedure repeated twice with 200 µL extraction solvent. Biological triplicates were performed and analyzed by HPLC-MS/MS. Measured values were mathematically converted into cellular c-di-GMP concentration. Constants of *E. coli* cell volume and cfu/mL needed for calculation were experimentally determined. A standard curve correlating *pdeH* induction and c-di-GMP levels was used to interpolate c-di-GMP values below the detection limit of around 10 nM c-di-GMP. To establish precise intracellular c-di-GMP levels, an *E. coli* Δ*pdeH* mutant was engineered with a plasmid-encoded IPTG inducible copy of *pdeH* (P_*lac*_::*pdeH*, pAR81). Engineered strains also harboring different reporter constructs were grown overnight in the presence of 1 mM IPTG or without IPTG. After diluting precultures into fresh medium containing variable concentrations of IPTG, cells were grown to an OD_600_ of 0.5, harvested and lysed for c-di-GMP measurements.

### Absolute protein concentration determination via selected reaction-monitoring (SRM) LC-MS analysis

600 µL of *E. coli* cultures grown in TB to an OD_600_ of ca. 0.5 were pelleted and washed twice with 1 mL ice-cold PBS. Cells were lysed in 1 % sodium deoxycholate (SDC), 100 mM Tris pH 8.5 by sonication. Proteins were denatured by heating at 95°C for 10 min. Protein alkylation was performed with chloracetamide. Protein digestion was performed by subsequent treatment with Lys-C (enzyme/protein ratio 1:200) and trypsin (enzyme/protein ratio 1:50). Peptides were acidified with TFA and desalted using PreOmics (ThermoFisher) cartridges. An aliquot of heavy reference peptide mix was spiked into each sample at a concentration of 20 fmol of heavy reference peptides per 1 µg of total endogenous protein mass. The heavy peptide mix contained 10 chemically synthesized proteotypic peptides (JPT Peptide Technologies GmbH) of the two target proteins (5 peptides each) that showed the highest MS1 responses in a previous large-scale study (Schmidt et al., 2016). In a first step, selected reaction-monitoring (SRM) assays (Peterson et al., 2012) were generated from a mixture containing 500 fmol of each reference peptide and shotgun LC-MS/MS analysis on a Q-Exactive HF platform. The setup of the μRPLC-MS system was as described previously (Ahrné et al., 2016).

Chromatographic separation of peptides was carried out using an EASY nano-LC 1000 system (Thermo Fisher Scientific), equipped with a heated RP-HPLC column (75 μm × 37 cm) packed in-house with 1.9 μm C18 resin (Reprosil-AQ Pur, Dr. Maisch). Peptides were analyzed per LC-MS/MS run using a linear gradient ranging from 95% solvent A (0.1% formic acid) and 5% solvent B (99.9% acetonitrile, 0.1% formic acid) to 45% solvent B over 60 minutes at a flow rate of 200 nL/min. Mass spectrometry analysis was performed on Q-Exactive HF mass spectrometer equipped with a nanoelectrospray ion source (both Thermo Fisher Scientific). Each MS1 scan was followed by high-collision-dissociation (HCD) of the 10 most abundant precursor ions with dynamic exclusion for 20 seconds. Total cycle time was approximately 1 second. For MS1, 3e6 ions were accumulated in the Orbitrap cell over a maximum time of 100 ms and scanned at a resolution of 120,000 FWHM (at 200 m/z). MS2 scans were acquired at a target setting of 1e5 ions, accumulation time of 50 ms, and a resolution of 30,000 FWHM (at 200 m/z). Singly charged ions and ions with unassigned charge state were excluded from triggering MS2 events. The normalized collision energy was set to 27%, the mass isolation window was set to 1.4 m/z and one microscan was acquired for each spectrum.

The acquired raw files were searched against a decoy database using the MaxQuant software (Version 1.0.13.13) containing standard and reverse sequences of the predicted SwissProt entries of *E. coli* (www.ebi.ac.uk, release date 2016/05/02), retention time standard peptides, and commonly observed contaminants (in total 10402 sequences) generated using the SequenceReverser tool from the MaxQuant software (Version 1.0.13.13). The precursor ion tolerance was set to 10 ppm, and fragment ion tolerance was set to 0.02 Da. The search criteria were set as follows: full tryptic specificity was required (cleavage after lysine or arginine residues unless followed by proline), 3 missed cleavages were allowed, carbamidomethylation (C) was set as fixed modification and arginine (+10 Da), lysine (+8 Da) and oxidation (M) were set as a variable modification. The resulting msms.txt file was converted to a spectral library panel with the 5 to 10 best transitions for each peptide using an in-house software tool. This was then imported into the SpectroDive program (Version 7.5, Biognosys, Schlieren, Switzerland), and a transition list for quantitative SRM analysis was generated. Here, all samples were analyzed on a TSQ-Vantage triple-quadrupole mass spectrometer coupled to an Easy-nLC (Thermo Fisher, Scientific). In each injection, an equivalent of 1.5 μg of peptides including heavy peptide references was loaded onto a custom-made main column (Reprosil C18 AQ, 3 μm diameter, 100 Å pore, 0.75 × 300 mm) and separated using the same gradient mentioned above. The mass spectrometer was operated in the positive ion mode using ESI with a capillary temperature of 275 °C, a spray voltage of +2200 V. All of the measurements were performed in an unscheduled mode and a cycle time of 2 sec. A 0.7 FWHM resolution window for both Q1 and Q3 was set for parent- and product-ion isolation. Fragmentation of parent-ions was performed in Q2 at 1.2 mTorr, using collision energies calculated with the SpectroDive software (version 7.5). Each condition was analyzed in biological quadruplicates. All raw files were imported into SpectroDive for absolute peptide and protein quantification.

### Biofilm assays

Attachment assays were carried out as described previously (Boehm et al., 2009). Briefly: 200 µL TB medium provided in a clear 96-well microtiter plate were inoculated 1:40 with an o/n culture grown at 37°C. The plate was incubated statically at 37°C for 24 h unless indicated differently. After recording the OD_600_ of the total biomass, the planktonic phase of the culture was discarded and the wells washed with H_2_O_dest._ from a hose. The remaining attached biomass was stained with 200 µL 0.3 % crystal violet (0.3 % (w/v) in 5 % (v/v) 2- propanol, 5 % (v/v) methanol) for 20 min. The plate was washed with H_2_O_dest._ from a hose and the stained biofilm dissolved in 20 % acetic acid for 20 min. The intensity of crystal violet stain was quantified at 600 nm and normalized to the initially measured total biomass.

For biofilm escape assays, cells harboring pAR81 were allowed to attach to plastic surfaces in 96-well microtiter plates for 7 hrs at 30°C in TB medium supplemented with 30 µg/ml ampicillin. Plates were gently washed with deionized water, dried, and incubated for 3 h at 30°C after adding fresh TB media was supplemented with IPTG to induce plasmid-borne P_lac_-*pdeH* and. After incubation, 10 µL of the planktonic phase in the center of the well were isolated to determine cfu/mL by spotting serial dilutions in LA plates supplemented with ampicillin. A pipetting robot (Tecan freedom evo) that allows for sensing of the liquid surface was used to improve reproducibility.

### Bacteriophage N4 propagation and infection assay

Phage N4 was propagated on *E. coli* MG1655 (CGSC 6300) and stored at 4°C in SM(G) buffer containing 0.1 M NaCl, 10 mM MgSO_4_ and 0.05 M Tris-HCl pH 7.5. Phage titer was determined by spotting 2.5 µl of a 10- fold serial dilution on a lawn of *E. coli* MG1655 via top agar overlay method as described before (Kropinski et al., 2009) with slight modifications. In brief, 100 µl of *E. coli* overnight culture was mixed with 3 ml top agar (0.4% LB-agar supplemented with 20 mM MgSO_4_ and 5 mM CaCl_2_) and pipetted on an LB agar plate pre-warmed to 60°C. After the top agar had solidified, 2.5 µl drops of serial dilutions of phage N4 (in PBS) were spotted on top and dried into the agar surface. Plates were incubated at 37°C overnight before N4 plaques were counted and compared between host strains.

### Flow cytometry

For flow cytometry of TB cultures, TB medium supplemented with 100 µg/ml ampicillin, 30 µg/ml kanamycin, and 0 or 65 µM IPTG was inoculated from single colonies. For strains carrying the c-di-GMP sensor, the medium was additionally supplemented with 200 nM anhydrotetracycline (aTc). TB cultures were grown overnight at 37°C and shaking. The next day cultures were diluted 1:500 in fresh TB medium supplemented with 40 µg/ml ampicillin, 20 µg/ml kanamycin, various IPTG concentrations, and where necessary, 200 nM aTc and grown at 37°C for 4.5 h. At this point, cell density was between OD_600_ 0.3-0.5 for all cultures. Cultures were diluted 1:200 in fresh TB medium with identical supplements and grown at 37°C for another 1.5 to 3.5 h up to a total incubation time of 6 to 8 h. Samples of max 800 µl were taken, kept on ice and diluted into 1x PBS just before analysis.

Flow cytometry of M9 minimal medium cultures started with day cultures in LB supplemented with 100 µg/ml ampicillin, which were inoculated from single colonies and grown for 6-7 h at 37°C. Cells were washed once in M9 medium without carbon source and diluted to an OD_600_ of 0.01 or lower in 5 ml M9 medium supplemented with 0.4% glycerol, 0.4% glycerol and 0.05% casamino acids, 0.5% glucose, 0.2% fumarate or 0.2% α- ketoglutarate. M9 cultures were incubated at 37°C with shaking at 180 rpm (Multitron, INFORS HT), for at least 17 h before measurements. At this point, cell density was between OD_600_ 0.1-0.4 for all cultures. To ensure exponentially growing cultures at the moment of sampling, M9 overnight cultures were diluted 1:500 in fresh M9 minimal medium with identical carbon source once in between where necessary. Samples were kept on ice and diluted into 1x PBS just before analysis.

Cells were analyzed on a BD LSRFortessa flow cytometer at medium flow rate and a maximum event rate of 10’000 events/s. Per sample, 100’000 events were recorded. Parameters measured were forward scatter (FSC-H), and side scatter (SSC-H and SSC-W). GFP was excited at 488/20 and detected with a 512/20 emission filter. Where applicable, mScarlet-I was excited using the pre-set excitation/emission ‘mCherry’. Data was collected using the Diva (BD Biosciences) software. FlowJo™ Software (version 10.6.1 for Windows) was used for the import and gating of raw data. The forward-scatter (FSC-H) and side-scatter (SSC-H and SSC-W) were used to separate cells from background particles. For analysis of cultures carrying the c-di-GMP sensor, a third gate was applied in which we used the ‘mCherry’ channel to gate for cells expressing mScarlet-I. Only mScarlet-I-positive cells were included in the analysis of the GFP signal coming from the c-di-GMP sensor. Gated populations were exported as ‘Scale Values’ in csv-files, and GFP distributions were visualized using MATLAB version R2019b (MathWorks) scripts.

## Author contributions

Conceptualization, A.R., B.S., F.F., M.v.B., A.H., T.S. and U.J.; Methodology, A.R., B.S., F.F., M.v.B., A.K., S.O., P.M., M.S.; Formal Analysis, A.R., B.S., F.F., M.v.B., A.K., S.O., P.M., M.S., T.S. and U.J.; Investigation, A.R., B.S., F.F., M.v.B., A.K., S.O., P.M., M.S., T.S. and U.J.; Resources, U.J. and T.S.; Writing – original draft, A.R., T.S and U.J. with contributions from all other authors; Funding acquisition, T.S. and U.J.

## Acknowledgements

We thank Tim Sharpe (Biophysics Facility, Biozentrum, University of Basel) for technical guidance and Fabienne Hamburger for cloning and strain construction. We thank Prof. Volkhard Kaever from the Medizinische Hochschule Hannover for c-di-GMP measurements. Moreover, we thank the Proteomics Core Facility (PCF) of the Biozentrum, University of Basel for absolute protein concentration quantification. This work was supported by the Swiss National Science Foundation (310030B_185372) to U.J., Swiss National Science Foundation (31003A_166652) to T.S. and Swiss National Science Foundation (PP00P3_170607) to C.P. The authors declare no competing interests.

## Data availability

The Protein Data Bank accession numbers for the coordinates of the structure reported here is 7PK5.

