## Supplementary Data for "Digital control of c-di-GMP in *E. coli* balances population-wide developmental transitions and phage sensitivity"

### Supplementary Figures

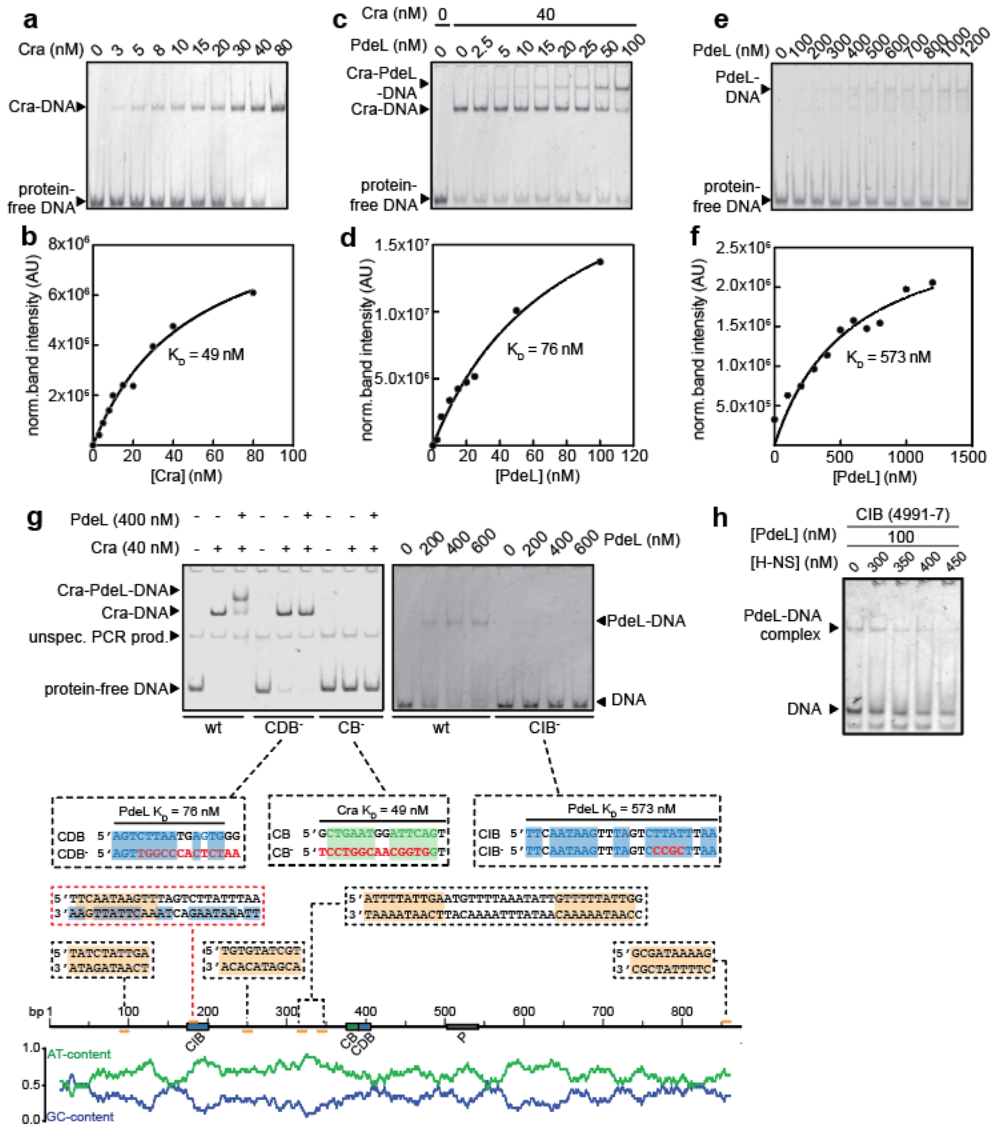


**Figure S1: Binding specificity and affinities of Cra and PdeL to *pdeL* promoter region. (a)** Binding of purified Cra-StrepII to the *pdeL* intergenic region as tested by electrophoretic mobility shift assay (EMSA). Binding was assayed using 10 nM of Cy3-labeled oligonucleotide spanning the Cra-box (CB) and 10 nM of Cy3-labeled oligonucleotide spanning the Cra-dependent PdeL-box (CDB) (see: Fig. 1a). Cra concentrations used are indicated. **(b)** Saturation binding fit of band intensities from (a). **(c)** EMSA and **(d)** binding affinity of purified PdeL-StrepII in the presence of 40 nM Cra-StrepII using the same oligos as in (a). The PdeL binding constant was calculated from band intensities of the super-shift (Cra-PdeL-DNA-complex). **(e)** Binding of purified PdeL-StrepII to the Cra-independent PdeL-box (CIB). The labeled oligonucleotide included the CIB region and 10 bp up- and 33 bp downstream of CIB. **(f)** Saturation binding fit of band intensities from (e). **(g)** Binding specificity of Cra and PdeL to the *pdeL* promoter region using oligos with mutated binding sites. Left panel: binding of Cra (40 nM) and PdeL (400 nM) to wild type (CB, CDB) and scrambled (CDB$^-$, CB$^-$) binding sites, respectively. Right panel: binding of PdeL (200 - 600 nM) to wild type and scrambled (CIB$^-$) CIB binding site. Sequences below the graphs indicate Cra and PdeL binding sites with scrambled sequences indicated in red in the lower strain. Note that mutations abolishing binding of PdeL to CDB or CIB were chosen outside of the Cra and H-NS consensus sequences, respectively. The bottom part of the graph shows putative H-NS binding boxes within the *pdeL* promoter region. PdeL and Cra binding sites are indicated in blue and green, respectively. H-NS binding sites (orange) were identified using the Virtual Footprint website http://www.prodoric.de/vfp/ (Münch et al., 2005). H-NS binding to CIB (red box) was experimentally verified in (h). AT- and GC-content of *pdeL* intergenic region with a binning of 5 bp is shown at the bottom. **(h)** PdeL and H-NS compete for CIB binding. EMSA assay with labeled DNA covering CIB (see Fig. 1a) and concentrations of purified proteins as indicated.

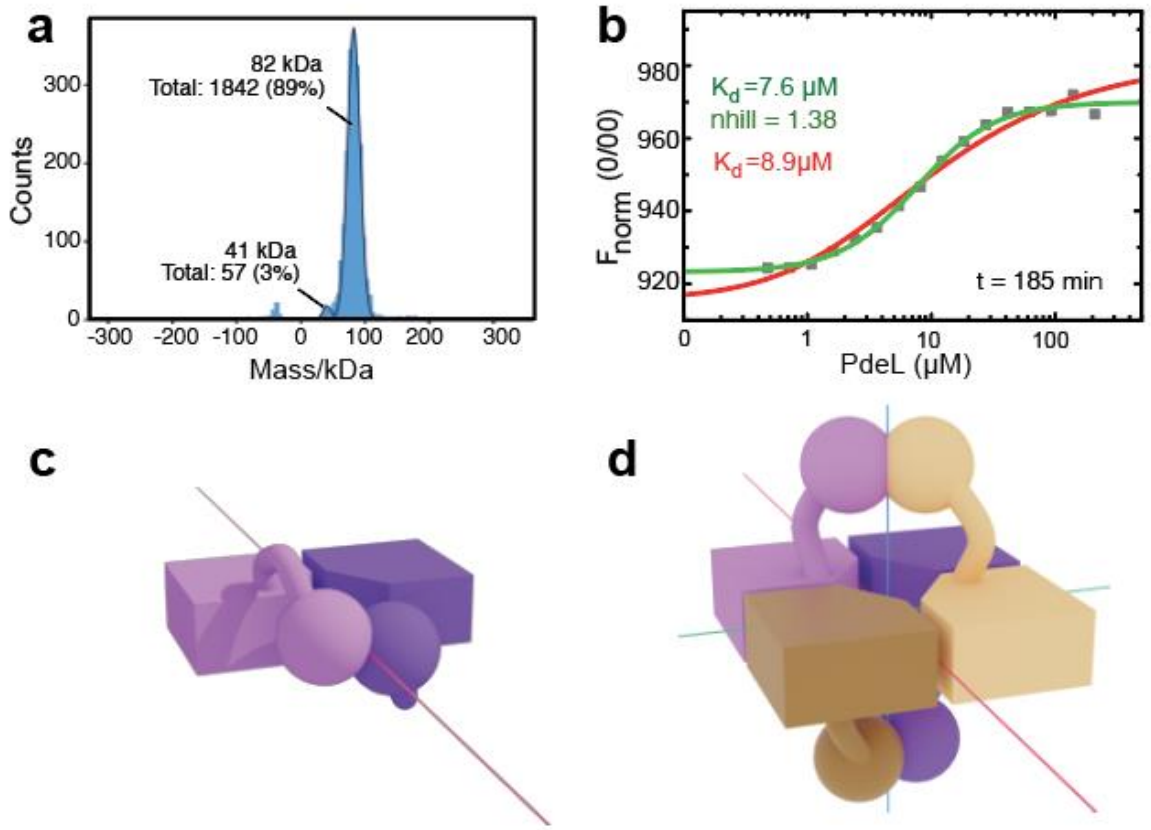


**<u>Figure S2</u>: PdeL oligomerization. (a)** PdeL is a dimer at a low protein concentration. Mass photometry analysis with mass histograms shown for 1899 steps of 25 nM PdeL. A Gaussian model is shown in black and the distribution of monomer (41 kDa) and dimers (82 kDa) are indicated. **(b)** MST signal of PdeL concentration series after equilibration (t = 185 min) with theoretical fit curve of a dimerization of dimers model (red), as shown in Fig. 2b. The fit of an allosteric sigmoidal curve is shown in green. **(c)** Schematic model of the R-state dimer of full-length PdeL. The EAL (prism) and HTH (sphere) domains of one subunit are associated with their counterparts by 2-fold symmetric interactions (red symmetry axis). **(d)** Schematic model of the R-state tetramer of full-length PdeL. The representation is as in (c), with a second dimer shown in brown hues and additional symmetry axes shown in blue and green. Note that for steric reasons only HTH domains of diagonally arranged EAL domains and not of neighboring EAL domains can meet to form canonical HTH dimers.

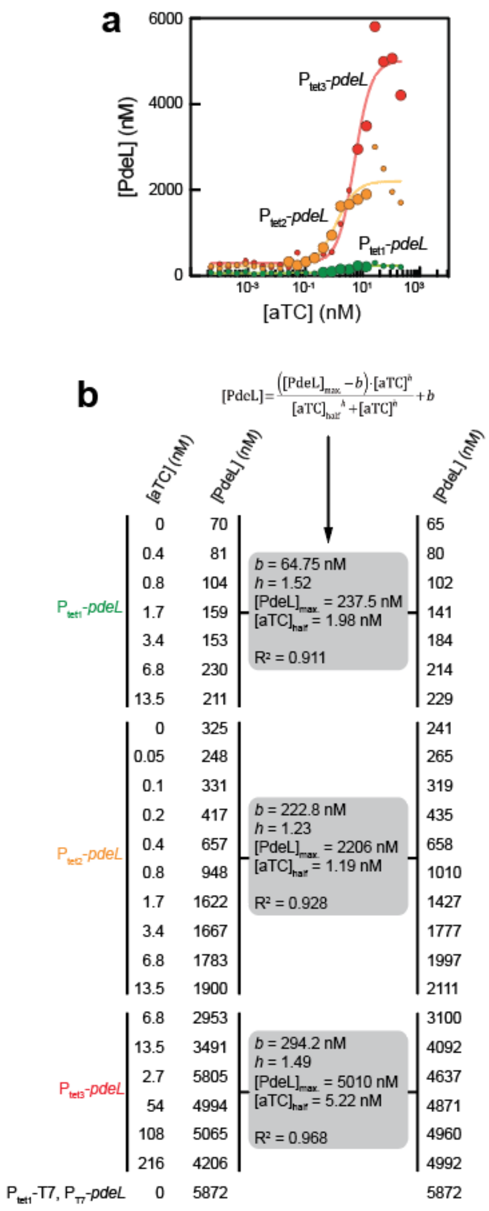


**<u>Figure S3</u>: Tuning of cellular levels of PdeL. (a)** PdeL protein concentration as measured by SRM (see Material and Methods). PdeL expression was tuned with three different $P_{tet}$-*pdeL* constructs harboring weak (green), intermediate (orange), and strong RBS (red) sequences providing different translational activities. For very high *pdeL* expression, a plasmid-based construct was used, in which $P_{tet1}$-driven T7 polymerase drives the expression of *pdeL* under control of the T7 promoter. The large dots depict conditions chosen to analyze the activity of the *pdeL* promoter over a broad range of PdeL concentrations (0.65 – 5.9 µM). **(b)** Data points of each $P_{tet}$ construct used in (a) were fitted with a allosteric sigmoidal curve (see Materials and Methods). The quality of individual fits is indicated with $R^2 > 0.9$. Smoothened PdeL concentrations were calculated from fit parameters.

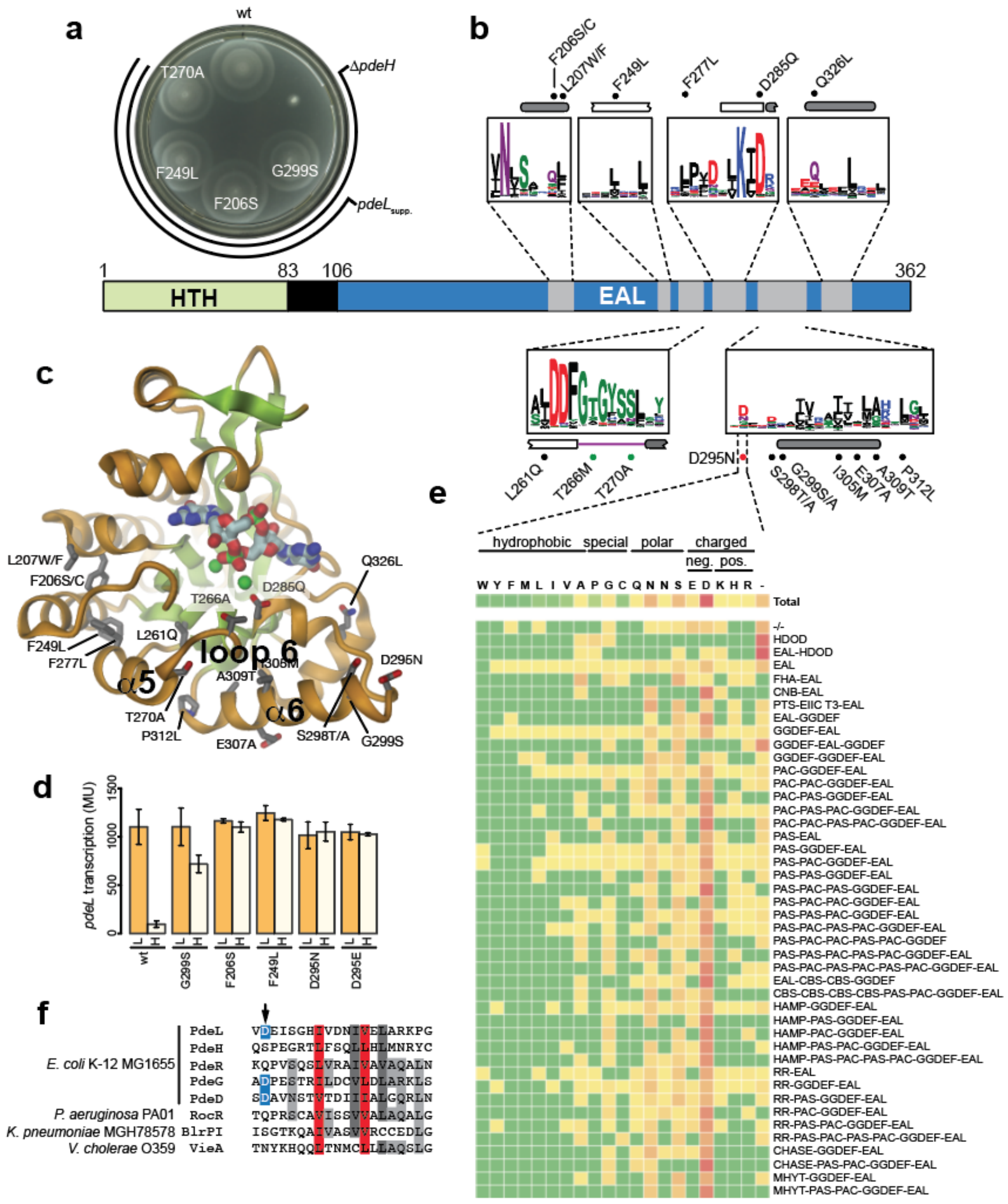


**Figure S4: Location and properties of activating *pdeL* alleles. (a)** Motility plate with *pdeL* suppressors restoring motility in a Δ*pdeH* mutant. **(b)** Domain architecture of PdeL with the DNA binding domain (HTH) in green and the catalytic EAL domain in blue. Suppressor mutations in the highly conserved loop 6 (green dots), in the R-state stabilizer Asp295 (red dot), or elsewhere in the *pdeL* coding sequence (black dots) are indicated. Alpha-helices (rounded grey bars), β-sheets (blank rectangles) and unstructured regions (line) are marked. Conservation of regions containing motile suppressors is shown as WebLogos of an alignment of 500 non-redundant EAL-domain proteins. **(c)** Crystal structure of PdeL_EAL in its T-state conformation with $Ca^{2+}$ (green dots), c-di-GMP (sticks) and positions of suppressor mutations (sticks and labels). **(d)** Activity of a *pdeL-lacZ* transcriptional reporter introduced into a selection of *pdeL* suppressor strains from (b) and (c). Low (L) and high (H) levels of c-di-GMP were established as indicated in Fig. 1f. **(e)** Conservation scores of the aspartic acid residue corresponding to D295 of PdeL in 500 non-redundant EAL-domain proteins with different domain architectures (see Material and Methods). Green and red colors indicate low and high occurrence, respectively. Amino acids are classified according to their chemical properties. EAL = phosphodiesterase; GGDEF = diguanylate cyclase. **(f)** Alignment of the PdeL region containing D295 with a selection of PDEs. All PDEs of *E. coli* K-12 with a conserved Asp (blue) at this position are listed.

**a**

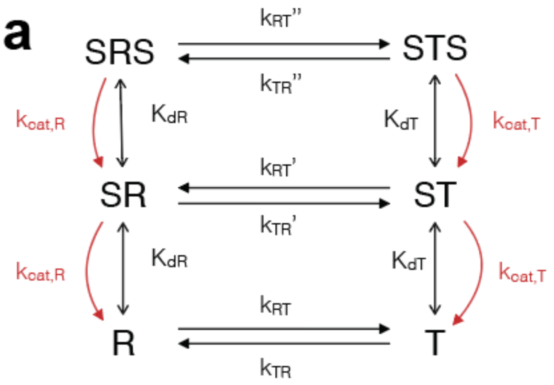


**b**

| | |
|---|---|
| $K_{d,R}$ [μM] *) | 1.0 |
| $K_{d,T}$ [μM] *) | 0.1 |
| $k_{cat,R}$ [$s^{-1}$] | 0.30 |
| $k_{cat,T}$ [$s^{-1}$] | 0.022 |
| $K:=k_{RT}/k_{TR}$ | 5.6 |
| $k_{RT}$ [$s^{-1}$] | $1.0 \cdot 10^{-3}$ |

*): fixed

**<u>Figure S5</u>: Kinetic model for the regulation of PdeL catalysis. (a)** Kinetic model describing the dynamic equilibrium between PdeL dimers in R- and T-conformation and their singly (RS, TS) or doubly (SRS, STS) occupied states, and the irreversible catalytic transitions (shown in red, with catalytic products not shown). The model assumes that there is no substrate cooperativity for enzyme binding. Note that the equilibrium constant K:=kRT /kTR is equal to T/R. **(b)** Parameters obtained from the fit to PdeL wild-type progress curves (Fig. 3e). Kd,R and Kd,T were chosen to account for an assumed 10-fold tighter binding of the substrate to the T- than to the R-state and to be consistent with the observed sub-micromolar affinity of the substrate to PdeL. Note that the fit was sensitive to the difference in the Kd values, but not to their exact absolute values, because the data were acquired at high substrate concentration (100 μM). For simplification, kRT' and kRT'' were set equal to kRT. Consequently, kTR' and kTR'' are given by kTR*KdT/KdR and kTR*(KdT/KdR)2, respectively, as dictated by pertinent thermodynamic cycles.

**Table S1. Data collection and refinement statistics.**

| | |
|---|---|
| **Resolution range (Å)** | 24.94 - 4.4 (4.556 - 4.4) |
| **Space group** | P 1 21 1 |
| **Unit cell (Å, deg.)** | 86.408 170.209 93.085 90 94.374 90 |
| **Total reflections** | 33559 (3369) |
| **Unique reflections** | 16827 (1691) |
| **Multiplicity** | 2.0 (2.0) |
| **Completeness (%)** | 98.20 (98.88) |
| **Mean I/sigma(I)** | 7.81 (1.81) |
| **Wilson B-factor ($Å^2$)** | 174.06 |
| **R-merge** | 0.05227 (0.3847) |
| **R-meas** | 0.07392 (0.5441) |
| **R-pim** | 0.05227 (0.3847) |
| **CC1/2** | 0.999 (0.86) |
| **Reflections used in refinement** | 16827 (1681) |
| **Reflections used for R-free** | 837 (105) |
| **R-work** | 0.3731 (0.3749) |
| **R-free** | 0.3972 (0.3557) |
| **CC(work)** | 0.816 (0.753) |
| **CC(free)** | 0.644 (0.714) |
| **Number of non-hydrogen atoms** | 7956 |
| **Number of molecules/a.u.** | 4 |

| | |
|---|---|
| **Protein residues** | 1012 |
| **RMS(bonds) (Å)** | 0.014 |
| **RMS(angles) (deg.)** | 1.48 |
| **Ramachandran favored (%)** | 96.41 |
| **Ramachandran allowed (%)** | 3.59 |
| **Ramachandran outliers (%)** | 0.00 |
| **Rotamer outliers (%)** | 6.88 |
| **Clashscore** | 5.35 |
| **Average B-factor (Å$^2$)** | 112.15 |
| **PDB-code** | 7PK5 |

Statistics for the highest-resolution shell are shown in parentheses.

**Table S2: Strains**

| Strain | Genotype | Source/Reference |
|---|---|---|
| CGSC6300 | E. coli K-12 MG1655 | Coli Genetic Stock Center (Yale) |
| CGSC7740 | E. coli K-12 MG1655 | Coli Genetic Stock Center (Yale) |
| BL21 (AI) | $F^-$ *ompT hsdS*$_B$ ($r_B^-$ $m_B^-$) *gal dcm araB*::T7 RNAP-*tetA* | Life techn. |
| AB330 | λ cI857 Δ*(cro-bioA)* | (Boehm et al., 2010) |
| AB607 | Δ*pdeH*::frt | (Boehm et al., 2010) |
| AB989 | λ cI857 Δ*(cro-bioA) kan*::$P_{rha}$-*ccdB-lacZ* | (Boehm et al., 2010) |
| AB2137 | Δ*pdeH*::frt *pdeL (G299S)-3xflag*::frt | (Reinders et al., 2015) |
| AB2202 | Δ*pdeH*::frt *pdeL (F206S)-3xflag*::frt | (Reinders et al., 2015) |
| AB2203 | Δ*pdeH*::frt *pdeL (F249L)-3xflag*::frt | this study |
| AB2271 | Δ*pdeH*::frt Δ*pdeL*::frt | this study |
| AB2377 | Δ*pdeH*::frt *kan*::$P_{pdeL}$-*lacZ* (merodiploid translational fusion) | this study |
| AB2378 | Δ*pdeH*::frt *pdeL (G299S)-3xflag*::frt, kan::$P_{pdeL}$-lacZ (merodiploid translational fusion) | this study |
| AB2400 | *kan*::$P_{pdeL\ CB-}$-*lacZ* (merodiploid translational fusion) | this study |
| AB2401 | *kan*::$P_{pdeL\ CB-\ \&\ CDB-}$-*lacZ* (merodiploid translational fusion) | this study |
| AB2402 | *kan*::$P_{pdeL\ CDB-}$-*lacZ* (merodiploid translational fusion) | this study |
| AB2519 | *pdeL (E235A)-3xflag*::frt *kan*::$P_{pdeL}$-*lacZ* (merodiploid translational fusion) | this study |
| AB2520 | *pdeL (D263N)-3xflag*::frt *kan*::$P_{pdeL}$-*lacZ* (merodiploid translational fusion) | this study |
| AB2521 | *pdeL (S298F)-3xflag*::frt *kan*::$P_{pdeL}$-*lacZ* (merodiploid translational fusion) | this study |
| AB2535 | *pdeL (G299S)-3xflag*::frt *kan*::$P_{pdeL}$-*lacZ* (merodiploid translational fusion) | this study |
| AB2569 | Δ*pdeL*::frt *kan*::$P_{pdeL}$-*lacZ* (merodiploid translational fusion) | (Reinders et al., 2015) |
| AB2571 | Δ*pdeH::frt* Δ*pdeL::frt* | this study |
| AB2609 | *pdeL (K60A)-3xflag*::frt *kan*::$P_{pdeL}$-*lacZ* (merodiploid translational fusion) | (Reinders et al., 2015) |

| | | |
|---|---|---|
| AB2727 | *kan*::$P_{pdeL\ P1}$-*lacZ* (merodiploid translational fusion) | this study |
| AB2731 | *pdeL (E141A)-3xflag*::frt *kan*::$P_{pdeL}$-*lacZ* (merodiploid translational fusion) | this study |
| AB2789 | Δ*pdeL::frt* Δ*cra*::frt *kan*::$P_{pdeL}$-*lacZ* (merodiploid translational fusion) | this study |
| AB2806 | Δ*cra*::frt *kan*::$P_{pdeL}$-*lacZ* (merodiploid translational fusion) | this study |
| AB2830 | Δ*hns*::frt *pdeL-3xflag*::frt *kan*::$P_{pdeL}$-*lacZ* (merodiploid translational fusion) | this study |
| AB2846 | Δ*pdeH*::frt *pdeL (D263N)-3xflag*::frt, kan::$P_{pdeL}$-lacZ (merodiploid translational fusion) | this study |
| AB2847 | Δ*pdeH*::frt *pdeL (K60A)-3xflag*::frt, kan::$P_{pdeL}$-lacZ (merodiploid translational fusion) | this study |
| AB2848 | Δ*pdeH*::frt *pdeL (S298F)-3xflag*::frt, kan::$P_{pdeL}$-lacZ (merodiploid translational fusion) | this study |
| AB2849 | Δ*pdeH*::frt *pdeL (E141A)-3xflag*::frt, kan::$P_{pdeL}$-lacZ (merodiploid translational fusion) | this study |
| AB2851 | Δ*pdeH*::frt *pdeL (E235A)-3xflag*::frt, kan::$P_{pdeL}$-lacZ (merodiploid translational fusion) | this study |
| AB2905 | *pdeL (K283R)-3xflag*::frt *kan*::$P_{pdeL}$-*lacZ* (merodiploid translational fusion) | this study |
| AB2907 | Δ*pdeH*::frt *pdeL (K283R)-3xflag*::frt, kan::$P_{pdeL}$-lacZ (merodiploid translational fusion) | this study |
| AB2923 | Δ*pdeH*::frt Δ*hns*::frt *pdeL-3xflag*::frt, kan::$P_{pdeL}$-lacZ (merodiploid translational fusion) | this study |
| AB2937 | *pdeL (D263N) (K283R)-3xflag*::frt, kan::$P_{pdeL}$-lacZ (merodiploid translational fusion) | this study |
| AB2939 | Δ*pdeH*::frt *pdeL (D263N) (K283R)-3xflag*::frt, kan::$P_{pdeL}$-lacZ (merodiploid translational fusion) | this study |
| AB2940 | *pdeL (F206S)-3xflag*::frt *kan*::$P_{pdeL}$-*lacZ* (merodiploid translational fusion) | this study |
| AB2942 | Δ*pdeH*::frt *pdeL (F206S)-3xflag*::frt, kan::$P_{pdeL}$-lacZ (merodiploid translational fusion) | this study |
| AB2943 | *pdeL (F249L)-3xflag*::frt *kan*::$P_{pdeL}$-*lacZ* (merodiploid translational fusion) | this study |
| AB2945 | Δ*pdeH*::frt *pdeL (F249L)-3xflag*::frt, kan::$P_{pdeL}$-lacZ (merodiploid translational fusion) | this study |

| | | |
|---|---|---|
| AB2986 | *kan*::$P_{pdeL}$-*lacZ* (merodiploid translational fusion) | this study |
| AB2996 | Δ*pdeH*::frt *csrA*::Tn5Δ(*kan*)::frt | this study |
| AB2997 | Δ*pdeH*::frt Δ*pdeL:*:frt *csrA*::Tn5Δ(*kan*)::frt | this study |
| AB3292 | *kan*::$P_{pdeL\ CIB-}$-*lacZ* (merodiploid translational fusion) | this study |
| AB3299 | Δ*pdeH*::frt frt::$P_{const.\ weak.}$-*pgaA-D* | this study |
| AB330 | λ cI857 Δ*(cro-bioA)* | this study |
| AB3302 | Δ*pdeH*::frt Δ*pdeL*::frt frt::$P_{const.\ weak.}$-*pgaA-D* | this study |
| AB3335 | Δ*pdeH*::frt $P_{pdeL\ CIB-}$-*pdeL-3xflag*::frt, kan::$P_{pdeL}$ CIB--lacZ (merodiploid translational fusion) | this study |
| AB3340 | Δ*pdeH*::frt *kan*::$P_{pdeL\ CDB-}$-*lacZ* (merodiploid translational fusion) | this study |
| AB3368 | Δ*pdeH*::frt *pdeL (T270A)-3xflag*::frt | this study |
| AB3381 | *pdeL (T270A)-3xflag*::frt *kan*::$P_{pdeL}$-*lacZ* (merodiploid translational fusion) | this study |
| AB3383 | Δ*pdeH*::frt *pdeL (T270A)-3xflag*::frt, kan::$P_{pdeL}$-lacZ (merodiploid translati-onal fusion) | this study |
| AB3431 | Δ*pdeH*::frt *pdeL*-[$RBS_{synth.}$-*mCherry*]$_2$::frt | this study |
| AB3447 | *pdeL (D295N)-3xflag*::frt *kan*::$P_{pdeL}$-*lacZ* (merodiploid translational fusion) | this study |
| AB3449 | Δ*pdeH*::frt *pdeL (D295N)-3xflag*::frt, kan::$P_{pdeL}$-lacZ (merodiploid translati-onal fusion) | this study |
| AB3455 | Δ*pdeH*::frt frt::$P_{tet}$-*tetR-pdeL*, kan::$P_{pdeL}$-lacZ (merodiploid translational fu-sion) | this study |
| AB3496 | Δ*pdeH*::frt *pdeL (D263N) (K283R)*-[$RBS_{synth.}$-*mCherry*]$_2$::frt | this study |
| AB3499 | Δ*pdeH*::frt $P_{pdeL\ CIB-}$-*pdeL*-[$RBS_{synth.}$-*mCherry*]$_2$::frt | this study |
| AB3501 | *$P_{pdeL}$ (CIB-)-pdeL::frt* | this study |
| AB3508 | Δ*pdeH*::frt frt::$P_{tet}$-*tetR-pdeL*, kan::$P_{pdeL}$ CIB--lacZ (merodiploid translatio-nal fusion) | this study |
| AB3673 | *pdeL (E141A)::frt* | this study |
| AB3718 | *pdeL (D295N)::frt* | this study |

| | | |
|---|---|---|
| AB3871 | *ΔpdeH::frt frt::Ptet-RBS (synth. strong)-tetR-RBS (synth. weak)-pdeL (L168R)* | this study |
| AB4490 | *CGSC 6300 ΔpdeH::frt* | this study |
| AB4491 | *CGSC 6300 ΔpdeL::frt* | this study |
| AB4513 | *pdeL (E141A)::frt-kan-frt* | this study |
| AB4514 | *pdeL (D295N)::frt-kan-frt* | this study |
| AB4519 | *CGSC 6300 pdeL (E141A)::frt* | this study |
| AB4520 | *CGSC 6300 pdeL (D295N)::frt* | this study |
| AB4532 | $P_{pdeL}$ *(CIB-)-pdeL::frt-kan-frt* | this study |
| AB4533 | *ΔpdeH::frt frt::Ptet-RBS (synth. strong)-tetR-RBS (synth. weak)-pdeL (L168R)::Frt-kan-Frt* | this study |
| AB4534 | CGSC 6300 Δ*(*$P_{pdeL}$*-pdeL)::frt-cat-frt* | this study |
| AB4536 | CGSC 6300 $P_{pdeL}$ *(CIB-)-pdeL::frt* | this study |
| AB4540 | CGSC 6300 *pdeL (L186R)::frt* | this study |

**Table S3: Plasmids**

| Plasmid | Genotype | Source |
|---|---|---|
| pCP20 | $FLP^{+}$ (*amp*) | (Cherepanov and Wackernagel, 1995) |
| pET28a | pBR332 lacI $P_{T7}$ (*kan*) 6xHis expression vector | Novagen |
| pKD46 | λ $RED^{+}$ (*amp*) | (Datsenko and Wanner, 2000) |
| pNDM220 | *repA parR parM* $P_{lac}$ (amp) | (Gotfredsen and Gerdes, 1998) |
| pAR1 | $P_{lac}$-*pdeL-strepII* in pET28a (*kan*) | (Reinders et al., 2015) |
| pAR3 | $P_{lac}$-*pdeL (D263N)-strepII* in pET28a (*kan*) | this study |
| pAR19 | $P_{lac}$-*cra-strepII* in pET28a (*kan*) | this study |
| pAR28 | $P_{lac}$-*hns-strepII* in pET28a (*kan*) | this study |
| pAR52 | $P_{lac}$-*pdeL (K283R)-strepII* in pET28a (*kan*) | this study |
| pAR62 | $P_{lac}$-*pdeL (D263N) (K283R)-strepII* in pET28a (*kan*) | this study |
| pAR81 | $P_{lac}$-$RBS_{synth.}$-*pdeH-3xflag* in pNDM220 (*amp*) | this study |
| pAR201 | $P_{lac}$-*pdeL (D295N)-strepII* in pET28a (*kan*) | this study |
| pAR202 | $P_{lac}$-*pdeL*$_{EAL}$*-3xflag-strepII* in pET28a (*kan*) | this study |
| pAR205 | $P_{lac}$-*pdeL*$_{EAL}$ *(Y268C)-3xflag-strepII* in pET28a (*kan*) | this study |
| pAR210 | $P_{lac}$-*pdeL*$_{EAL}$ *(Y268C) (T270A)-3xflag-strepII* in pET28a (kan) | this study |
| pAR211 | $P_{lac}$-*pdeL*$_{EAL}$ *(Y268C) (E235A)-3xflag-strepII* in pET28a (kan) | this study |
| pAR212 | $P_{lac}$-*pdeL*$_{EAL}$ *(Y268C) (D295N)-3xflag-strepII* in pET28a (kan) | this study |
| pAR226 | *pdeL-3xFlag::kan* in pUC19 *(amp)* | this study |

| | | |
|---|---|---|
| pAR231 | *pdeL::kan* in pUC19 (*amp*) | this study |
| pAR323 | PpdeL-gfpmut2 in pUA66 (kan) | this study |
| pAR341 | $P_{lac}$-RBSsynth.-pdeH-3xflag in pNDM220k (kan) | this study |
| p2H12ref | $P_{tet}$-sensor-mScarlet-I (ampicillin) | UJ11206, Kaczmarczyk & Jenal, unpublished |
| p2H12ref-blind | $P_{tet}$-sensor*-mScarlet-I (ampicillin) | UJ11207, Kaczmarczyk & Jenal, unpublished |

#### Table S4: Oligos

| Primer # | Description | Restriction Site | Sequence |
|---|---|---|---|
| 4991 | CIB fwd | | Cy3-GTTGCGAATGTTCAATAAGTTTAG |
| 4997 | CIB rev | | Cy3-ATGCGTCATTTCAAATGATCAGC |
| 4695 | CB CDB fwd | | Cy3-TGCTGAATGGATTCAGTCTTAATGAGTGGG |
| 4696 | CB CDB rev | | Cy3-CCCACTCATTAAGACTGAATCCATTCAGCA |
| 4283 | RNAP interaction | | Cy3-GAGCAAAGGCGCATTATATG |
| 4398 | Construction of AB2986/AB2727/AB2569/AB2400/AB2401/AB3292 | | TCGCTTGCTGTCTCCGGAACTAG-TCGAGGCCATGGTGGCCTTGTTATTGCATAAAACCGCGCC |
| 4399 | Construction of AB2986/AB2727/AB2569/AB2400/AB2401/AB3292 | | CGTTGTAAAACGACGGCCAGTGAATCCG-TAATCATGGTCATCAGAAAAACAC-GAAAATCACATGAATTCATGAACACACCTTTATCTTTTATC |
| 5032 | AB2727 | | GCTATGTGTTTATGAGGGAGAGAGCGCGCTGGTTGATCTG-GATTAGCCCGCCCATATAATGCGCCTTTGCTCATG |
| 5033 | AB2727 | | CTCTCCCTCATAAACACATAGC |
| 5028 | AB2402 | | CATTTGCTGAATGGATTCAGTTGGCCCAC-TCTAATTTTTAAGGGACAGGCATAGAG |
| 5029 | AB2402 | | ACTGAATCCATTCAGCAAATG |
| 5030 | AB2400 | | GGCATTGCTATAATATTGGTTATCATTTTCCTGGCAAC-GGTGGTCTTAATGAGTGGGTTTTTAAG |
| 5031 | AB2400 | | AAATGATAACCAATATTATAGCAATGCC |
| 5036 | AB2401 | | GGCATTGCTATAATATTGGTTATCATTTTCCTGGCAAC-GGTGGTTGGCCCAC-TCTAATTTTTAAGGGACAGGCATAGAG |
| 5037 | AB2401 | | AAATGATAACCAATATTATAGCAATGCC |
| 5248 | AB3292 | | CATTTGCTGAATGGATTCAGTCATTATGAG-TGGGTTTTTAAGGGACAGGCATAGAG |
| 5249 | AB3292 | | CACGAAAATCACATGAATTCATGAACACAC-CTTTATCTTTTATC |
| 14546 | AB4490 | | CGCAGCGGTTATTTTGCCTGCCTTG |
| 14547 | AB4490 | | GAGGAAATTTACCCCGGCCAGGCGG |
| 1675 | AB4491/AB4519/AB4520 | | AATGGCATATTAACGGGGATCTATCAGG |
| 4125 | AB4491/AB4519/AB4520 | | GCGTAACTGTTTTTGAGCGAG |

| | | | |
|---|---|---|---|
| 12053 | AB4513/AB4514/AB4533/AB4536/AB4540 | | CGCTGCCAACCTCCCGATGCGCTGTGAATAGCGG |
| 13550 | AB4513/AB4514/AB4533/AB4536/AB4540 | | GGTAATAAATTTATCTCTGAATGG |
| 10528 | AB3718 | | CATTTGTGCAAATGGCGAG-TGTCAATGAAATCTCCGGTCATATTGTGGAC |
| 10529 | AB3718 | | GTCCACAATATGACCGGAGATTTCATTGACAC-TCGCCATTTGCACAAATG |
| 16111 | AB4540 | | GACAGGTAATCACTCATACTGAACAGCGATAAAAGA-TAAAGGTGTGTTCATGAATTCATGTGATTTTCGTGTTTTTCTGC |
| 16112 | AB4534 | | CAGATAATAAAAAAGAAAAGAACTATTGCAGCCCAAAAC-CTACATTT-GGGCTGAGCGATTGTGTAGGCTGGAGCTGCTTC |
| 16113 | AB4534 | | CGGGAGAATTAATCGCTGCCAAC-CTCCCGATGCGCTGTGAATAGCGGTTTACAGAA-TATCCTCCTTAGTTCCTATTCCG |
| 4126 | AB4536 | | GGCCATAGCGTGATGTTCC |
| 4697 | pAR1/pAR3/pAR52/pAR62/pAR201 | NcoI/NotI | CAAAAAAACCATGGCTAATTCATGTGATTTTCGTG |
| 4698 | pAR1/pAR3/pAR52/pAR62/pAR201 | NcoI/NotI | CAAAAAAA-GCGGCCGCTTACTTTTCGAACTGCGGGTGGCTCCAACCAC-CTGCTTTCATTACCCATTC |
| 4403 | pAR19 | NcoI/XhoI | CAAAAACCATGGCAAAACTGGATGAAATCGCTCGGCTG |
| 4404 | pAR19 | NcoI/XhoI | CGTTTTTCTCGAGTTACTTTTCGAACTGCGGGTGGCTCCA |
| 5839 | pAR28 | NcoI/NotI | CAAAAAAACCATGGCTAGCGAAGCACTTAAAATTCTG |
| 5840 | pAR28 | NcoI/NotI | CAAAAAAA-GCGGCCGCTTACTTTTCGAACTGCGGGTGGCTCCATT-GCTTGATCAGGAAATCGTCG |
| 7323 | pAR81 | BamHI/XhoI | GAAAAAAAGGATCCATAGGAGGAACAATTTTATGA-TAAGGCAGGTTATCCAGC |
| 7324 | pAR81 | BamHI/XhoI | GAAAAAAACTCGAGTTATTTATCGTCGTCATCTTTGTAG-TCGA-TATCATGATCTTTATAATCACCGTCATGGTCTTTGTAGTCGAATAGCGCCAGAACCGCCGTATTC |
| 8812 | pAR202 | NcoI/NotI | GAAAAAAA-GCGGCCGCTTACTTTTCGAACTGCGGGTGGCTCCATTTATCGTCGTCATCTTTGTAGTCGATATC |
| 10524 | pAR202 | NcoI/NotI | GAAAAAAACCATGGCTGATAATCACTATCACCATATCG |
| 8271 | pAR205 | NcoI/NotI | GCGCTGGATGACTTTGGTACGGGTTGTGCGAC-CTATCGTTACTTGCAGG |
| 8272 | pAR205 | NcoI/NotI | CCTGCAAGTAACGATAGGTCGCACAACCCGTAC-CAAAGTCATCCAGCGC |
| 8812 | pAR205/pAR210/pAR211/pAR212 | NcoI/NotI | GAAAAAAA-GCGGCCGCTTACTTTTCGAACTGCGGGTGGCTCCATTTATCGTCGTCATCTTTGTAGTCGATATC |
| 10524 | pAR205/pAR210/pAR211/pAR212 | NcoI/NotI | GAAAAAAACCATGGCTGATAATCACTATCACCATATCG |
| 10639 | pAR210 | NcoI/NotI | CTGGATGACTTTGGTACGGGTTGTGCGGCC-TATCGTTACTTGCAGGCG |
| 10640 | pAR210 | NcoI/NotI | CGCCTGCAAGTAACGATAGGCCGCACAACCCGTAC-CAAAGTCATCCAG |

| | | | |
|---|---|---|---|
| 5150 | pAR211 | NcoI/NotI | CGTTACCGGAATAGGGTTACGTGCCGTTAGCTCGAGAAC-CAGCTTG |
| 5151 | pAR211 | NcoI/NotI | CGTAACCCTATTCCGGTAACG |
| 10528 | pAR201/pAR212 | NcoI/NotI | CATTTGTGCAAATGGCGAG-TGTCAATGAAATCTCCGGTCATATTGTGGAC |
| 10529 | pAR201/pAR212 | NcoI/NotI | GTCCACAATATGACCGGAGATTTCATTGACAC-TCGCCATTTGCACAAATG |
| 5163 | pAR3 | NcoI/NotI | CAACATTACCTTTGCGCTGGATAACTTTGGTAC-GGGTTATGCGACCTATC |
| 5164 | pAR3 | NcoI/NotI | ATCCAGCGCAAAGGTAATGTTG |
| 6424 | pAR52 | NcoI/NotI | GGCGTTCCCGGTCGATTTTATTCGGATCGATAAGTCATTT-GTGCAAATG |
| 6425 | pAR52 | NcoI/NotI | CATTTGCACAAATGACTTATCGATCCGAATAAAATCGAC-CGGGAACGCC |
| 11825 | pAR62 | NcoI/NotI | CTTTGCGCTGGATAACTTTGGTACGGGTTATGCGAC-CTATCGTTACTT-GCAGGCGTTCCCGGTCGATTTTATTCGGATCGATAAGTC |
| 11826 | pAR62 | NcoI/NotI | GACTTATCGATCCGAATAAAATCGACCGGGAACGCCTG-CAAGTAACGATAGGTCG-CATAACCCGTACCAAAGTTATCCAGCGCAAAG |
